# Ribosome changes elicit non-canonical translation for chemosurvival in G0 leukemic cells

**DOI:** 10.1101/2021.12.07.471635

**Authors:** C. Datta, SS. Truesdell, SIA. Bukhari, H. Ngue, B. Buchanan, Keith Q. Wu, O. Le Tonqueze, S. Lee, M. Granovetter, M. Boukhali, J. Kreuzer, W. Haas, S. Vasudevan

**Author notes:** Correspondence & contact: Shobha Vasudevan 185 Cambridge St, CPZN4202, Boston MA 02114.

## Abstract

Quiescent leukemic cells survive chemotherapy, with translation changes. Our data reveal that FXR1, a protein amplified in several aggressive cancers, increases in quiescent and chemo-treated leukemic cells, and promotes chemosurvival. This suggests undiscovered roles for this RNA- and ribosome-associated protein in chemosurvival. FXR1 depletion decreases translation and ribosome subunits, with altered rRNAs, snoRNAs, and ribosomal proteins (RPs). We find that FXR1 binds factors that promote ribosome gene transcription and bind snoRNAs. Ribosome changes increased in FXR1-overexpressing cells, including increased snoRNAs and RPLP0/uL10, activate eIF2α kinases. Accordingly, phospho-eIF2α increases, enabling non-canonical translation of survival and immune regulators in FXR1-overexpressing cells. Overriding these with inhibitors reduces chemosurvival. Thus, increased FXR1 in quiescent or chemo-treated leukemic cells, alters ribosomes that trigger stress signals to re-direct translation for chemosurvival.

**One Sentence Summary:** FXR1 alters ribosomes in G0, which induce stress signals to elicit noncanonical translation for AML drug and immune survival.

## Main Text: Introduction

Cancer cells can enter a reversible arrest phase called quiescence or G0 that is resistant to harsh conditions including chemotherapy (*1–8*). We previously found that leukemic cells induced to G0 by growth factor deprivation, are not only chemoresistant, but also exhibit similar post-transcriptional changes, as that of leukemic cells surviving chemotherapy (*1*). This indicated translation of specific genes in such chemoresistant cells, which are needed for their chemosurvival. Understanding the altered translation program and how chemosurviving G0 cells translate specific genes, can reveal undiscovered insights on chemoresistance, and new strategies to reduce chemosurvival.

Canonical translation initiation is mediated by two rate limiting steps: cap dependent recruitment of mRNAs, and recruitment of the initiator tRNA/ternary complex for translation initiation (*9, 10*). Such conventional translation promotes proliferation associated genes (*11*). One of the two rate-limiting steps of translation initiation is inhibited by dephosphorylation of the canonical cap complex inhibitor, EIF4EBP (4EBP) due to low mTOR activity (*9, 12, 13*). The second rate limiting step of translation initiation is regulated by phosphorylation of the tRNA recruitment complex factor, eIF2α. Phosphorylation of eIF2α reduces its activity, which inhibits canonical translation. This is brought about by four eIF2α kinases (eIF2aks) that are triggered by various stress responses, and cause the integrated stress response (ISR) (*10, 14*). We found that G0 and chemotherapy-treated cells exhibit eIF2α phosphorylation (*1*), which can inhibit canonical translation, and enable specific genes to get translated non-canonically. How G0 and chemotherapy-treated cells induce eIF2α phosphorylation, to switch to non-canonical translation and express specific genes that lead to chemosurvival and AML persistence, remains to be uncovered.

Our previous data revealed that the RNA binding protein, Fragile-X-mental retardation related protein 1 (FXR1) (*15–25*) increases in serum-starved G0 acute monocytic leukemic (AML) THP1 cells. FXR1 has been shown to be important for tumor progression as it is amplified in several aggressive cancers, where post-transcriptional expression of specific mRNAs is altered (*26*). FXR1 is associated with ribosomes, translation, mRNA stability, and localization, and localizes in the nucleus, cytoplasm, and in stress granules (*15–25, 27*). In serum-starved G0 cells, we found that FXR1a splice isoform increases and can promote specific mRNA translation (*28, 29*). Given that FXR1 increases in serum-starved G0 AML cells and in aggressive cancers (*26*), and promotes specific translation in G0 AML cells (*28, 29*) that are chemoresistant (*1*), the role of FXR1 on chemosurvival via translation mechanisms needs to be uncovered, to understand the impact of translation regulation in chemosurviving cancer cells.

In this study, we investigated the changes in translation in G0 and chemosurviving AML cells, and the role of the enhanced FXR1 in G0 cells on AML chemosurvival. Our findings demonstrated that as in serum-starved G0 AML cells that are chemoresistant, FXR1 increases in therapy surviving AML cells. Consistently, our data reveal that cells overexpressing FXR1 show increased chemosurvival, while FXR1 depletion reduces chemosurvival. We find that the increased FXR1 associates with ribosome regulators and induces changes in ribosome components. These ribosomal changes trigger stress signaling via eIF2α kinase activation that causes eIF2α phosphorylation. This reduces canonical translation and permits non-canonical translation of specific pro-survival genes, leading to chemosurvival. Pharmacological inhibition of this induced non-canonical translation, or of the translated pro-survival genes, suppresses chemosurvival, indicating new avenues to therapeutically target refractory AML.

## Results

### FXR1 increases in Cytarabine-treated, chemosurviving cells

We recently showed that G0 cells, induced by serum-starvation, are chemoresistant with similar gene expression as in surviving cells isolated post-chemotherapy (*1*). We previously found that FXR1 increases in THP1 AML cells that are induced to G0 by serum deprivation (*28*). We found that both serum-starved G0 cells where FXR1 increases, and chemosurviving cells show similar translation factor changes compared to untreated, proliferating cells (Fig. 1Aa-b (*1*)) that may underlie their common ability to survive chemotherapy (*1*). Therefore, we asked the question whether FXR1, an RNA- and ribosome-associated, translation regulator, could be altered in levels and function in chemosurviving cells isolated after chemotherapy treatment. We tested FXR1 levels in Cytarabine (AraC) chemotherapy treated cells (*1*). We find that FXR1 increases in AraC-treated, surviving THP1 cells (AraC, Fig. 1Ac, S1A) and on serum-starvation (G0) (*28*) of THP1 cells that are chemoresistant (*1*). This was also observed upon AraC treatment or serum-starvation of another AML cell line (Fig. S1B), indicating that the increase of FXR1 upon AraC treatment was not unique to THP1 cells. These data reveal increase of FXR1 upon AraC therapy treatment in AML cells.

### THP1 cell survival, upon treatment with AraC chemotherapy, is promoted by FXR1 overexpression and reduced by FXR1 depletion

To investigate whether FXR1 plays a role in chemosurvival of G0 and AraC treated cells where it increases, we used a previously constructed THP1 cell line for FXR1 depletion. This is stably transduced with an inducible shRNA lentiviral vector that depletes all FXR1 isoforms (FXR1 KD) upon doxycycline induction, compared to a parallel control shRNA expressing cell line (control). These cells are induced with doxycycline to express the shRNA for three days to effectively deplete FXR1 (*28, 29*). We had previously seen that FXR1a isoform increases in G0 THP1 cells (*28*). Therefore, we also constructed a THP1 cell line that constitutively overexpresses FXR1a isoform (FXR1 OE) compared to a vector control cell line. Western blot analysis show that FXR1 was effectively depleted or overexpressed (Fig. 1B-C, Western blots). These were treated with AraC to test the impact of FXR1 levels on chemosurvival. Consistent with its increase in AraC-surviving cells, we find that FXR1 overexpression promotes chemosurvival by 1.9-fold while FXR1 depletion reduces chemosurvival to less than 50% (Fig. 1B-C graphs). These data indicate that FXR1 promotes chemosurvival of THP1 cells.

**Fig. 1.**
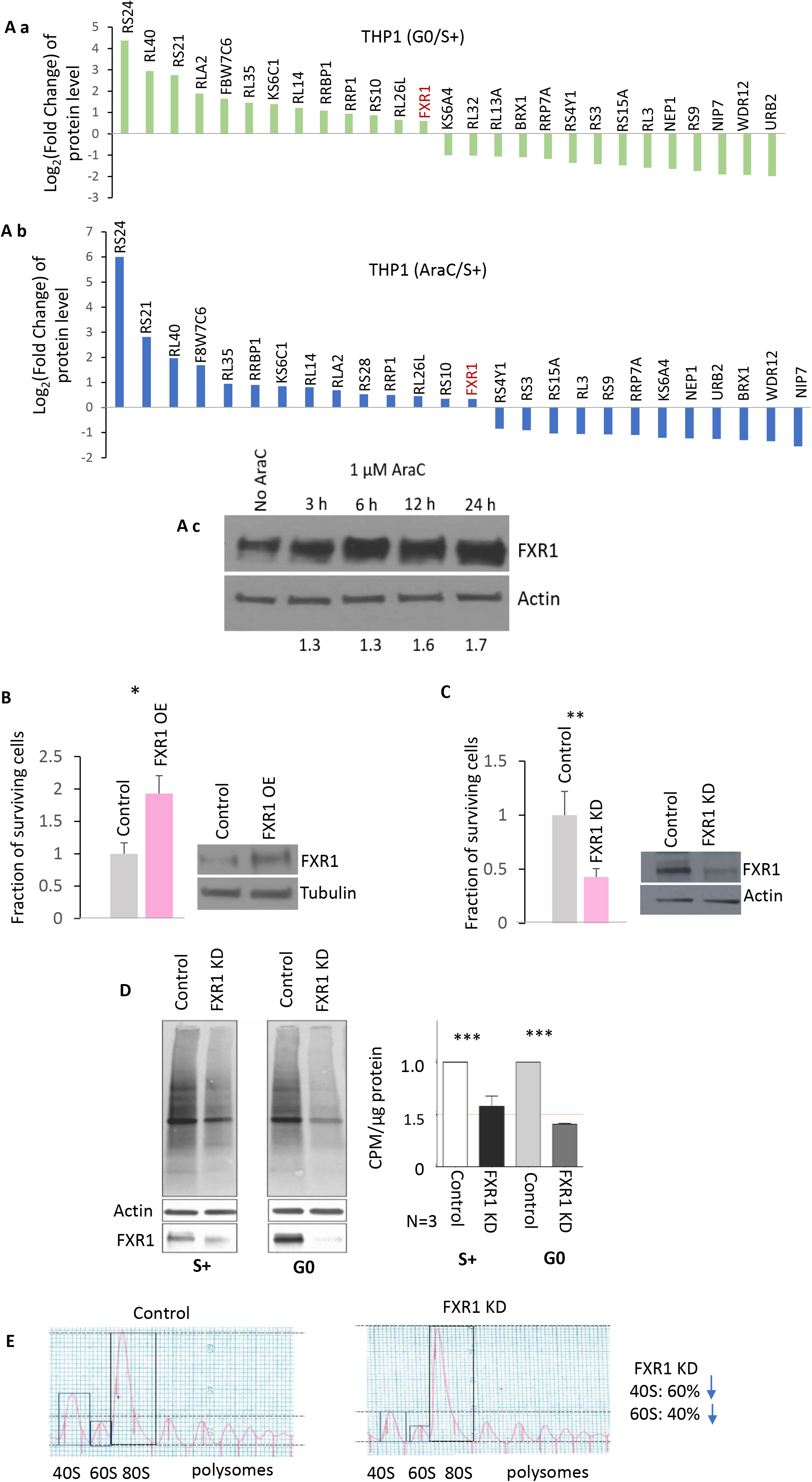
FXR1 is required for translation, and for 40S and 60S ribosome subunits levels. **A.** Graphs showing proteomic levels of commonly upregulated ribosomal proteins and regulators including the RNA- and ribosome-associated regulator, FXR1, in **a.** THP1 cells grown in serum starved medium (G0) and in **b.** THP1 cells treated with 5 µM AraC from (*1*). **c.** FXR1 levels over time of AraC (5µM) treatment compared to untreated cells. **B-C.** THP1 survival with AraC chemotherapy by trypan blue stain exclusion cell counts, **B.** with FXR1 overexpression (FXR1 OE) compared to vector control, and **C.** with FXR1 depletion compared to vector control shRNA cells; Western blot below of FXR1, with Tubulin and Actin as loading controls. Comparison of global translation in FXR1 knockdown (FXR1 KD) compared to shRNA control (Control) stable cell lines in untreated (S+) and serum-starved G0 cells by **D.** 35S Methionine labeling followed by SDS-PAGE; below Western analysis of FXR1 with Actin as loading control, and below quantification of 35S levels, and by **E.** polysome analyses in FXR1 KD and control G0 cells showing polysomes and ribosome subunits. Data are average of 3 replicates +/-SEM. See also Fig. S1.

### FXR1 depletion decreases overall translation

We next explored the mechanism of how FXR1 may promote chemosurvival. Our previous data showed that FXR1a overexpression and tethering to a reporter promoted overall translation (*28–30*). To investigate the effect of FXR1 on overall translation, we performed 35S Met incorporation analysis to label nascently translated proteins in FXR1 knockdown cells followed by SDS-PAGE separation and phosphorimager quantitation, or scintillation counter quantitation of the total levels of nascently labeled proteins. FXR1 depletion decreased the levels of protein synthesis as observed from the 35S Met incorporation by 30% in serum conditions and upto 50% decrease in serum-starved cells (Fig. 1D). These data suggest that FXR1 is needed for global translation.

### FXR1 depletion leads to decreased ribosome subunits

To test whether the significant decrease in translation was due to perturbations in ribosome levels, we conducted polysome analysis in FXR1 depleted cells compared to control shRNA cells (Fig. 1E). We find that both 60S and 40S subunits decreased significantly (40% and 60% respectively, Fig. 1E). These data indicate reduced ribosome subunits in FXR1 depleted cells.

### FXR1 depletion decreases rRNAs

Given the decrease in ribosome subunits, we investigated whether FXR1 levels alter ribosome biogenesis. Ribosome biogenesis includes rRNA transcription and processing, rRNA modification, and ribosomal proteins and their assembly. These involve several regulators that control steps from transcription to processing and assembly (*31–46*). Analysis of ribosomal RNAs (rRNAs) by qPCR revealed a significant decrease in 28S, 18S and 5.8S rRNA levels upon FXR1 depletion (Fig. 2A). These effects include both transcriptional and processing effects as 45S precursor rRNA, from which these mature rRNAs are derived, was also reduced upon FXR1 depletion. This decrease is not only because of Pol I transcription of the precursor of mature 28S, 18S, and 5.8S rRNAs, as the fourth rRNA, the 5S rRNA that is transcribed by Pol III, is also decreased (Fig. 2A). In contrast, FXR1 overexpression increased all rRNA levels (Fig. 2A), consistent with the previously noted increase in overall translation upon overexpression of FXR1(*30*). These data suggest that FXR1 depletion affects regulation of ribosome biogenesis.

**Fig. 2.**
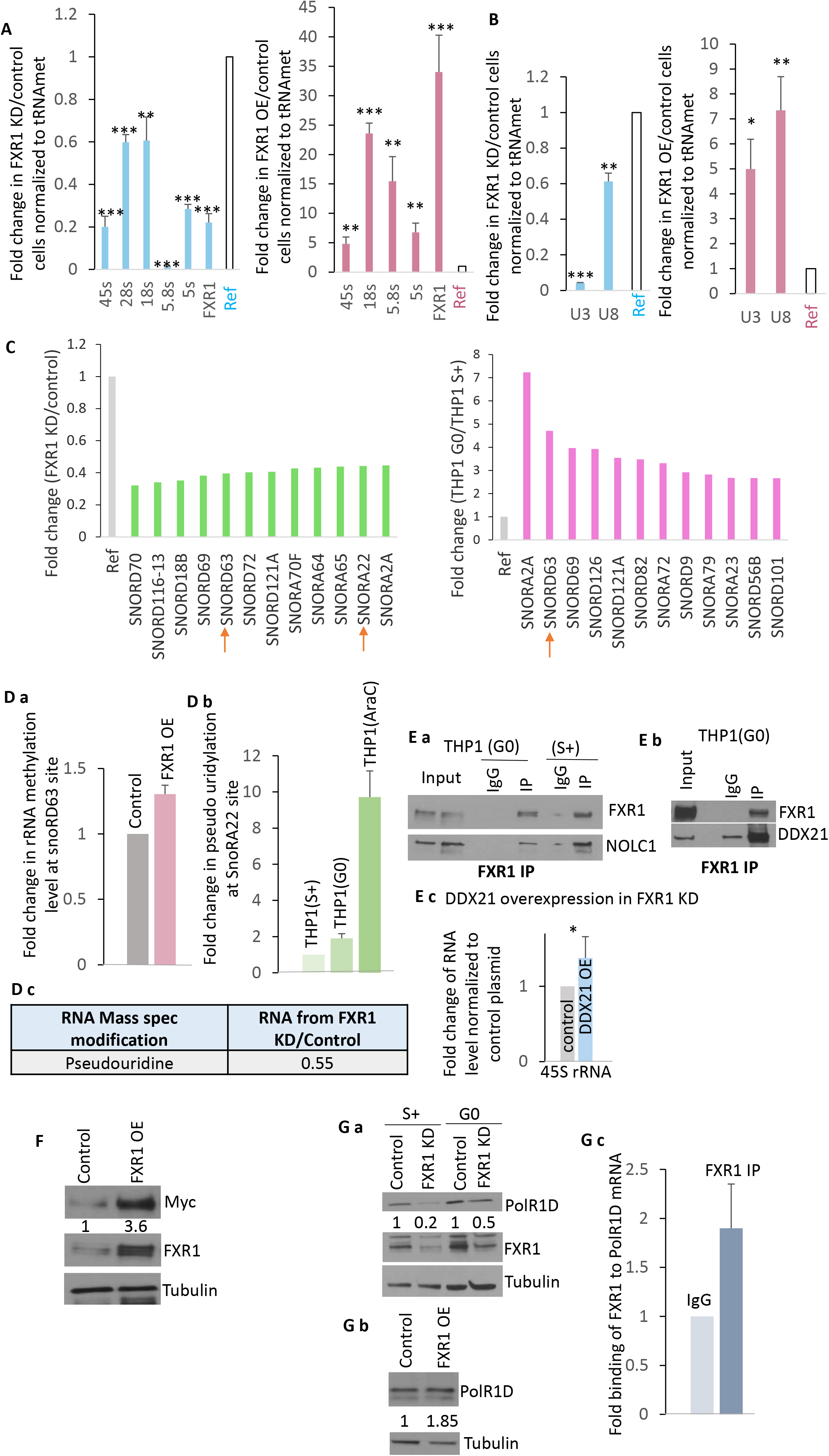
FXR1 regulates rRNA and snoRNA levels, and associates with ribosome and snoRNA regulators. **A.** 45S, 18S, 28S, 5.8S, & 5S RNAs, and **B.** snoRNAs that regulate rRNA cleavage and processing, U3 and U8, in FXR1 KD compared to control shRNA cells, and in FXR1 OE cells compared to control cells. Shown qPCR of rRNA and snoRNA levels normalized to tRNA-met. 28S values for FXR1 OE was larger than the scale. **C.** SnoRNA levels from microarray profiles in FXR1 KD G0 THP1 cells compared to control shRNA cells, and the levels of the same snoRNAs in THP1 G0 compared to serum-grown proliferating cells from Table S1a-d. **D. a.** 2′-*O*-methylation assay by low dNTP RT-qPCR, in FXR1 OE cells compared to control cells, of rRNA site (28 S rRNA A4541m) modified by a snoRD, snoRD63, that is regulated in G0 and by FXR1 from our dataset in Table S1a-d, and of **b.** rRNA modification site (on 28S at U4966 and U4975) of a snoRA, snoRA22, that is regulated in G0 and by FXR1 from the dataset in Table S1a-d, analyzed in THP1 (G0) and THP1 cells treated with AraC as measured by CMC treatment followed by low dNTP RT-qPCR assay. Mass Spectrometry data of **c.** pseudouridine in rRNA enriched samples from FXR1 KD compared to control G0 cells from Table S1g. **E.** FXR1 association with ribosome and snoRNA regulators, **a.** NOLC1 and **b.** DDX21, and others in Table S2. In vivo formaldehyde crosslinking coupled FXR1 immunoprecipitation followed by Western analysis of FXR1, NOLC1, and DDX21. **c.** Partial rescue of rRNA level (shown qPCR of 45S rRNA) by overexpression of DDX21 in FXR1 KD cells. **F.** Western blot of c-MYC levels in FXR1 OE compared to vector control cells. **G.** Western blot of POLR1D in **a.** FXR1 KD G0 and serum-grown (S+) proliferating cells, and **b.** in FXR1 OE compared to vector control cells. **c.** qPCR analysis for PolR1D mRNA in FXR1 immunoprecipitated RNA compared to IgG control immunoprecipitated RNA. Data are average of 3 replicates +/-SEM. See also Fig. S2, Tables S1and S2.

### SnoRNAs involved in rRNA processing and modification, increase in G0 or FXR1- overexpressing THP1 cells, and decrease upon FXR1 depletion

Ribosome biogenesis involves RNA regulators called small nucleolar RNAs (snoRNAs) that process or modify rRNAs. Specific snoRNAs involved in rRNA processing, such as U3 (snoRD3) and U8 (snoRD118), bind RNA-protein complexes (RNPs) that are required to cleave the 47S rRNA precursor for rRNA processing into mature 18S, 28S and 5.8S rRNAs. Many other snoRNAs bind RNA modification enzymes and other RNA binding proteins to form snoRNPs that are known to base pair and modify rRNAs at specific sites. Box C/D snoRNAs or snoRDs recruit Fibrillarin enzyme to cause 2′-*O*-methylation while Box H/ACA snoRNAs or snoRAs recruit Dyskerin enzyme to cause pseudouridylation of rRNA sites. These modifications affect ribosome interactions and functions (*33, 37, 47-56*).

We first examined levels of the snoRNAs involved in rRNA cleavage, U3 and U8. Global RNA profiling in G0 and proliferating cells revealed that U3 increases in G0 THP1 cells compared to proliferating, serum-grown cells (*1, 28, 29*) (Table S1a-d). Consistently, we find that U3 and U8 are decreased upon FXR1 depletion and increased upon FXR1 overexpression by qPCR analysis (Fig. 2B), which could impact rRNA levels in Fig. 2A. Global profiling also revealed that several snoRNAs that modify distinct rRNA sites also increase in G0 compared to proliferating THP1 cells (*1, 28, 29*) (Table S1a-d). We further find that many of the snoRNAs increased in G0 THP1 cells associate with FXR1 in global RNA profiling of FXR1 co-immunoprecipitates from in vivo crosslinked G0 THP1 cells (Table S1e-f). Consistently, we find that these snoRNAs that are increased in G0 THP1 cells where FXR1 increases, are decreased upon FXR1 depletion in global RNA profiles of FXR1 shRNA depleted cells compared to control shRNA THP1 cells (Fig. 2C, Table S1a-d). These data are consistent with the impact of FXR1 levels observed on ribosome levels and translation, and on rRNA levels (Fig. 1B-E, 2A).

### SnoRNAs regulated by FXR1 alter rRNA modification

To test the functional outcome of FXR1 interaction on snoRNAs, we analyzed specific rRNA sites in FXR1 depletion or overexpressed cells, as well as examined modifications on rRNAs globally in FXR1 depleted G0 cells compared to control G0 cells. To examine specific snoRNA target modification sites, we analyzed 2′-*O*-methylation by low dNTP RT-qPCR, as well as pseudouridylation after *N*-cyclohexyl-*N*′-(2-morpholinoethyl) carbodiimide metho-p-toluenesulfonate (CMC metho-*p*-toluene sulfonate)-treatment followed by low dNTP RT-qPCR (*55, 57–59*), at the rRNA sites of specific snoRNAs that are increased in G0 regulated by FXR1. To globally examine rRNA modification changes with or without FXR1, we enriched rRNA (devoid of poly(A) RNA and RNAs fractionated to remove RNAs less than 200nt including tRNAs) from FXR1 knockdown G0 cells and control cells and subjected the nucleosides to LC-MS analysis (Fig. S2B).

Our data show changes in modification at rRNA sites targeted by the snoRNAs that are increased in G0 and regulated by FXR1. As shown in Fig. 2Da, the snoRD63 target modification site on 28S rRNA (A4541m) shows enhanced 2′-*O*-methylation in FXR1 overexpression cells, consistent with snoRD63 increase in G0 cells where FXR1 increases, and its decrease in FXR1 knockdown cells. snoRA22 increases in G0, and we find that rRNA pseudouridylation at snoRA22 target sites on 28S rRNA (U4966 and U4975) increases in G0 and AraC cells (Fig. 2Db). Consistent with snoRA22 decrease upon FXR1 depletion, we find that FXR1 knockdown reduces overall pseudouridylation by mass spectrometry (Fig. 2Dc). Other known rRNA modifications also show changes in FXR1 knockdown cells by mass spectrometry (Fig. S2A, Table S1g). The sites targeted by many of these FXR1 regulated snoRNAs are key for tRNA interaction and translation (*60–66*). These data indicate that FXR1 alters levels and functions of specific snoRNAs that are increased in G0 cells, which can affect rRNA levels and modifications, to alter translation in these chemosurviving cells.

### FXR1 interacts with rRNA and snoRNA regulators, and modulates the levels of ribosome gene transcription factors

To identify how FXR1 may mediate effects on snoRNAs and on rRNA levels, we examined FXR1 immunoprecipitates for interacting regulators by in vivo crosslinking G0 THP1 cells followed by FXR1 immunoprecipitation and Tandem-Mass-Tag (TMT) spectrometric analysis (*28, 67*) of co-immunoprecipitates (Table S2). We find that FXR1 interacts with multiple ribosome and translation related proteins in G0 cells (Fig. S2Ba). We also find that in vivo crosslinking coupled FXR1 immunoprecipitation in control or FXR1 overexpression cells reveals FXR1 association with rRNAs (Fig. S2Bb). These data suggest that FXR1 may interact with and affect ribosome regulators.

Importantly, we find that snoRNA and ribosome regulators DDX21 and NOLC1, co-immuno- precipitated with FXR1 (Fig. 2Ea-b, S2Ba-b, Table S2), indicating that FXR1 may affect ribosome levels through such interactions with such ribosome regulators. DDX21 (*68, 69*) binds and affects the roles of snoRNAs that control rRNA modification and processing. DDX21 also promotes ribosome transcription via Pol I for rRNAs and Pol II for snoRNA and ribosomal protein expression. NOLC1 is a snoRNA and ribosome regulator that is involved in their assembly and biogenesis (*70–74*). Thus, regulators like DDX21 and NOLC1 are good candidates that could be mediating in part the effects of FXR1 on snoRNAs and rRNA transcription, and thereby, on translation. Consistently, overexpression of DDX21 in FXR1 KD cells, where rRNA levels decrease (Fig. 2A), partially rescued the rRNA defect by increasing levels of 45S rRNA (Fig. 2Ec). These data indicate that FXR1 levels may affect the functions of a key ribosome gene transcription and snoRNA regulator, DDX21, which can lead to the alterations in the ribosome observed.

FXR1 also associates with and regulates RNAs that control ribosome gene transcription and processing. This includes snoRNAs that co-immunoprecipitate with FXR1 in G0 cells (Table S1e-f). FXR1 has been recently shown to bind and regulate the 3′UTR of c-MYC mRNA (*75*), a major ribosome gene transcription regulator that affects all three polymerases and regulates ribosome biogenesis (*76, 77*). Consistently, we find that c-MYC is increased upon FXR1 overexpression (Fig. 2F), which could lead to increased ribosome biogenesis. Since FXR1 affects both Pol I transcript 45S rRNA and Pol III transcript 5S rRNA, we hypothesized that a common component of both Pol I and Pol III complexes may be affected by FXR1 and G0 chemoresistant cell conditions, leading to this coordinated impact on both 45S derived rRNAs and 5S rRNA in Fig. 2A. Therefore, we tested whether common Pol I and Pol III components are increased in G0 and are affected in FXR1 depleted or overexpressed cells. POLR1D is an essential component of both Pol I and Pol III complexes and is needed for DDX21 to locate to the nucleolus (*78*); thus, POLR1D is needed for both 45S and 5S rRNA production (*78*). Mass spectrometry dataset from G0 and AraC-treated cells reveal that POLR1D increases in G0 cells that also show FXR1 increase (Fig. S2C, Table S3a). Consistently, we find that POLR1D is reduced and overexpressed with FXR1 knockdown or overexpression respectively (Fig. 2Ga-b). Accordingly, we find that POLR1D mRNA associates with FXR1 (Fig. 2Gc), indicating that FXR1 associates with the mRNA of POLR1D to regulate levels of this rRNA transcription regulator. Thus, FXR1 regulation of the levels of a common factor of the Pol I and Pol III complexes, POLR1D, can affect levels of all rRNAs observed in Fig. 2A. Together, these data suggest that FXR1 interacts with or regulates the levels or functions of multiple ribosome biogenesis factors to alter ribosomes in G0 and AraC cells where FXR1 increases.

### G0 cells, AraC-treated cells, and FXR1 overexpressing cells alter ribosomal protein levels

Given that snoRNAs and rRNAs, and their regulators are associated with or modulated by FXR1 that increases in G0 and AraC-treated cells, and that ribosomal proteins and regulators are commonly altered in G0 and AraC-treated cells (Fig. 1Aa-b, Table S3a-b), we examined the other main ribosome component, ribosomal protein (RP) levels upon FXR1 depletion and overexpression. Many RPs modulate rRNA processing (*31, 35-38, 40, 43-46*) and RP genes are regulated by DDX21 and c-MYC that are modulated by FXR1 (Fig. 2E-F). Additionally, we find that some RPs interact with FXR1 in G0 cells (Fig. S2Ba, Table S2), consistent with previous studies that showed RPs interact with FXR1 (*79, 80*). Thus, RPs could be altered and could lead to changes in ribosome complexes (*81*) and translation output in G0 and AraC-treated cells where FXR1 increases.

We find that specific RPs increased or decreased in G0 are also increased in AraC treated cells in our proteomic and RNA analyses (Fig. 1Aa-b, Table S3b (*1*)), indicating that common changes in RP composition occur in G0 cells that are chemoresistant, and in AraC-surviving cells. Interestingly, we find many of these are decreased upon FXR1 depletion or increased in FXR1 overexpression cells by qPCR (Fig. 3A), and by RNA profiling analyses (Table S3c-e (*28, 29*)). Furthermore, RP changes were also observed in FXR1 depletion cells compared to control cells in G0, by polysome profiling (Table S3h-i), and by TMT-spectrometric proteome analysis (Table S3f-g). These include P stalk proteins RPLP0 and RPLP2, that are part of the GTPase activation center (GAC) that is needed for GTPase translation factors in translation elongation (*82*), as well as other RPs such as RPL29 that increase in G0 cells, in AraC treated cells, and in FXR1 overexpression cells, and conversely decrease in FXR1 depletion cells (Fig. 3Ba-c, S3C, Table S3). These data suggest differences in the ribosome in cells with increased FXR1 levels, as in G0 and AraC treated cells, which can impact ribosome complexes, and thereby alter ribosome function.

**Fig. 3.**
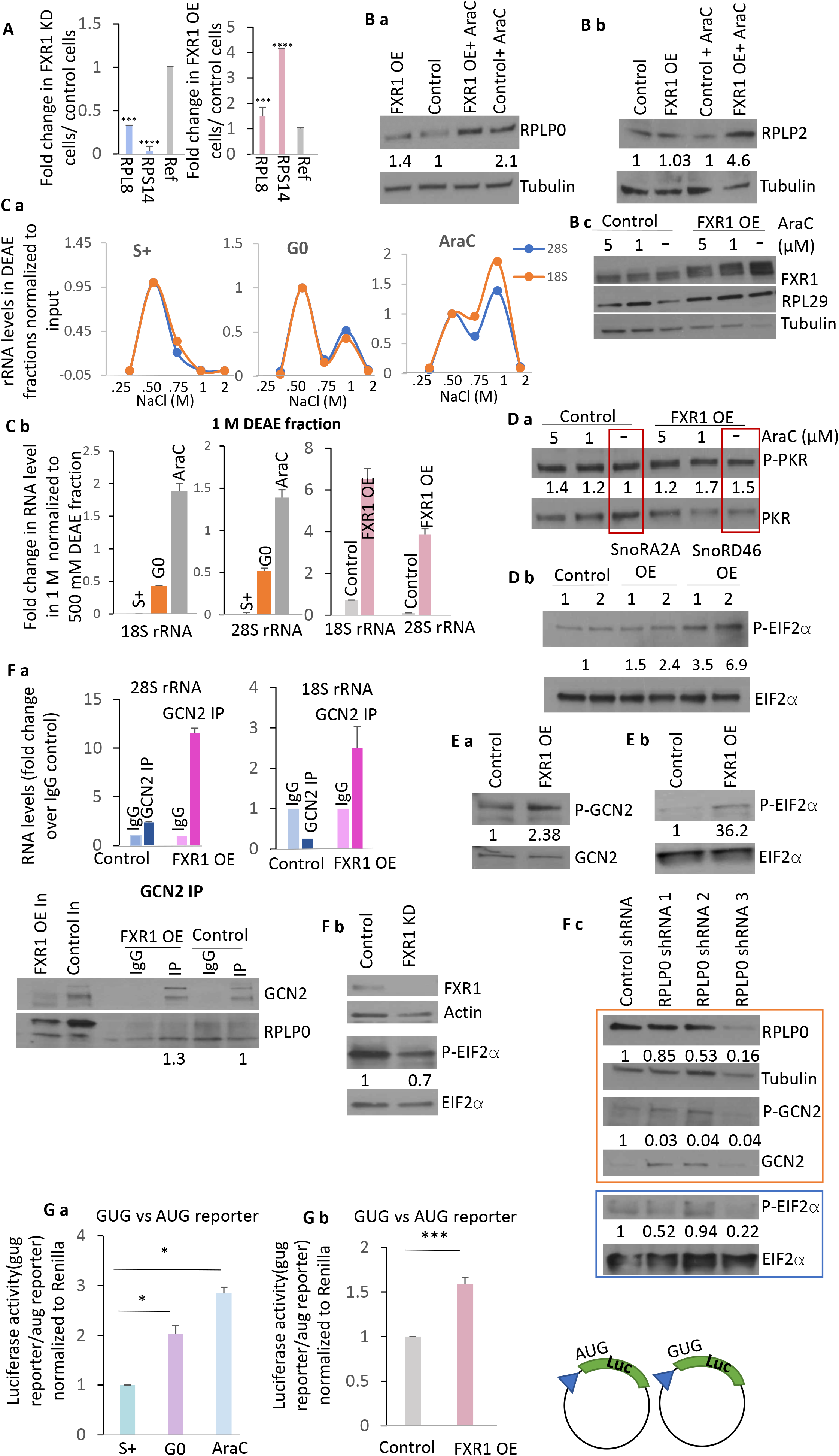
eIF2α phosphorylation and non-canonical translation increase with altered ribosomal components upon FXR1 overexpression, similar to G0 and AraC-treated cells. **A.** RP protein levels by qPCR normalized to tRNA-met in FXR1 KD cells compared to control shRNA cells, and in FXR1 OE cells compared to control cells from Table S3e. Western blot analysis of **B.** a. RPLP0, b. RPLP2, c. RPL29 in FXR1 OE and control cells with or without AraC treatment. **C.** a. Ribosome purification with Y10B immunoprecipitation from in vivo crosslinked cell cytoplasmic extracts, followed by DEAE fractionation to analyze ribosome complex migration in distinct salt fractions in G0 and AraC-treated cells compared to untreated S+ cells by qPCR analysis of fractions for where both 18S and 28S rRNAs (representing 80S ribosomes) co-migrate. b. DEAE fractionation and qPCR analysis of rRNAs of Y10B antibody immunopurified ribosome complexes (18S and 28S rRNAs) in the 1M fraction normalized for levels of the 500mM fraction, from G0 and AraC-treated cells compared to S+ cells, as well as from in vivo crosslinked FXR1 OE cells compared to control cells. **D.** Western blot of phosphorylation of a. of PKR in FXR1 OE cells compared to control cells, and with AraC treatment, normalized to total PKR protein levels. b. Overexpression of snoRA2A and snoRD46 (Fig. S3D for graph of qPCR showing snoRNA amplification) that cause increased rRNA 2′-*O*- methylation, also leads to increase of eIF2α phosphorylation as shown by Western analysis. **E.** Western blot of phosphorylation of a. GCN2 and b. eIF2α phosphorylation in FXR1 OE compared to control cells, normalized to their total protein levels. **F.** a. Immunoprecipitation of GCN2 followed by qPCR analysis of rRNA levels, and of RPLP0 and GCN2 by Western blot analysis (below), in FXR1 OE compared to control cells. b. Western blot analysis of eIF2α phosphorylation in FXR1 KD cells. c. Western blot analysis of GCN2 and eIF2α phosphorylation normalized to their total protein levels in FXR1 OE cells with control shRNA or RPLP0 depletion. **G.** Translation ratios of GUG start site Luciferase reporter over AUG reporter normalized to co-transfection Renilla control reporter, as indicated by Luciferase reporter assays (RNA levels in Fig. S3E) in a. G0 and AraC-treated cells compared to untreated, serum-grown THP1 cells, and in b. FXR1 overexpressing cells compared to control cells. Data are average of 3 replicates +/-SEM. See also Fig. S3, Table S3.

### Ribosome complexes in G0 or AraC-treated cells, migrate distinctly compared to proliferating, untreated cells, but similarly to ribosome complexes in FXR1 overexpressing cells

As rRNA processing, RP level, and modification changes are induced in G0 by FXR1 overexpression (Fig. 2, 3A-B), we analyzed ribosome complexes formed in AraC-treated and serum-starved G0 cells, to test if they are distinct compared to proliferating cells. We also compared these with ribosome complexes in FXR1 overexpressing cells compared to vector control cells, as they show similar chemosurvival and ribosome changes to G0 and AraC-treated cells. These cells were first subject to formaldehyde crosslinking to freeze in vivo complexes.

Cytoplasmic extracts were prepared to avoid nuclear, pre-ribosome complexes that are not assembled. To identify changes in ribosome complexes, the crosslinked cytoplasmic extracts were bound on an anionic column (DEAE) and eluted by increasing salt concentrations to separate complexes (*30, 83*). Fractions of the in vivo crosslinked complexes that contained both 18S *and* 28S rRNAs, depicting small and large subunits, were examined as composite 80S ribosome complexes. We observed one peak at 500 mM salt in proliferating, untreated, serum-grown cells; in contrast, we find a second distinct peak (1 M salt) of ribosome complexes in serum-starved G0 and AraC-treated cells (Fig. S3A). Strikingly, we observed two peaks, eluting in the same salt fractions (500mM, 1M), in FXR1 overexpressing cells compared to one (500mM) in vector control cells (Fig. S3A). These data indicate changes in ribosome complexes that could impact translation and chemosurvival that is commonly observed in G0, AraC-treated, and FXR1 overexpressing cells but not in untreated, proliferating control vector cells (Fig. 1-2). These data suggest that FXR1 overexpression in G0 and AraC cells may cause changes in ribosome complexes.

Next, we wanted to ensure that these complexes separated on DEAE are enriched for ribosomes. Therefore, we immunopurified ribosomes from G0 and AraC-treated cells compared to untreated, serum-grown cells, as well as FXR1 overexpression cells compared to vector control cells, and then examined them with DEAE fractionation. These cells were first subject to formaldehyde crosslinking to freeze in vivo ribosome complexes. Cytoplasmic extracts were prepared to avoid assembling, nuclear, pre-ribosome complexes. Ribosome complexes were purified from these cytoplasmic extracts by Y10B, an antibody that recognizes 5.8S complexes in assembled 80S complexes and ribosomes (Fig.S3Ba) (*83, 84*). Y10B immunopurification of assembled 80S ribosomes was verified by qPCR analysis of enrichment of 5.8S rRNA and

Western analysis of ribosomal protein RPLP0 in Y10B immunoprecipitates compared to IgG control (Fig. S3Bb). Lack of unprocessed 45S rRNA in Y10B immunoprecipitates verified the absence of partially processed pre-ribosomes in Y10B immunoprecipitates of 80S cytoplasmic ribosome complexes (Fig.S3Bb). To test whether Y10B-purified ribosome complexes migrate distinctly in FXR1 overexpressing cells, G0 cells, and AraC-treated cells compared to vector control cells or untreated, serum-grown (S+) cells, the Y10B immunoprecipitates were fractionated over DEAE and eluted with increasing salt. These were examined for fractions that showed both 18S and 28S rRNAs depicting small and large subunit rRNAs, as composite 80S ribosome complexes, by qPCR analysis. While untreated, serum-grown cells showed a single peak at 500 mM salt, a second complex is observed at 1M salt in G0 and AraC treated cells (Fig. 3Ca-b), consistent with the ribosome component changes observed in these conditions in Figs 2, 3A-B and in S3A. Interestingly, we see a similar pattern of two distinct peaks of ribosome complexes seen in G0 & AraC treated cells compared to untreated proliferating cells, eluting in the same salt fractions (500mM, 1M) in FXR1 overexpressing cells compared to the single lower salt peak in vector control cells (Fig. 3Cb). These data suggest that in FXR1 overexpression conditions that include artificially overexpressed cells, or serum-starved G0 cells and AraC-treated cells where FXR1 increases, ribosomal complex changes are observed. While this could reflect different ribonucleoproteins (RNPs) associated with the ribosome that cause differential complex migration, these data indicate that ribosomes are differentially bound or comprised in FXR1-overexpressing cells, and migrate in fractions that are also similarly observed in G0 and AraC-treated cells, but not in untreated, serum-grown proliferating cells.

### Altered P stalk proteins and snoRNAs in G0 cells, chemo-treated cells, and FXR1 overexpressing cells, increase eIF2α phosphorylation

Such multiple changes on the ribosome can alter many downstream mechanisms. One way that ribosomes can alter translation, is by activating stress signaling pathways (*85–90*). G0 and AraC surviving cells where FXR1 increases, show inhibition of canonical translation with increased phosphorylation of eIF2α (*1*). This is brought about by eIF2α kinases that respond to multiple stress signals, and can be induced by ribosomal component changes elicited by increased FXR1: via enhanced snoRNAs (*91–93*), and via ribosome changes in the RP P stalk proteins (*85–87, 94*).

FXR1 enhances levels of several snoRNAs (Table S1a-d) that have been shown to bind eIF2α kinase PKR (eIF2ak2), a dsRNA binding protein (*91–93*) that causes eIF2α phosphorylation. We hypothesized that increased snoRNAs by FXR1 amplification as in G0 and AraC surviving cells could, in part, lead to eIF2α phosphorylation, as snoRNAs have been shown to activate eIF2ak2 or PKR (*91–93*). Consistently, we find that FXR1 overexpressing cells and AraC-treated cells show increased phosphorylation and thus activation of PKR (Fig. 3Da). Therefore, we tested the impact on eIF2α phosphorylation upon overexpression of two snoRNAs that we found are enhanced in G0 and FXR1 overexpressing cells, snoRD46 and snoRA2A. We verified overexpression of these transfected snoRNAs by qPCR compared to a control vector (Fig. S3D). We find that overexpression of these snoRNAs leads to increased eIF2α phosphorylation, compared to expression of a control vector (Fig. 3Db). This indicates that overexpression of FXR1 increases snoRNAs, which can lead to activation of dsRNA binding eIF2α kinase, PKR (*91–93*) that can phosphorylate eIF2α.

Recent studies also show that P stalk proteins, RPLP2, RPLP1, and RPLP0 or uL10, promote phosphorylation of eIF2α by activation of GCN2 eIF2α kinase (eIF2ak4), via RPLP0 interaction (*85–87, 94*). Significantly, we find that RPLP0 increases in FXR1 overexpressing cells (Fig. 3Ca) and decreases upon FXR1 knockdown (Fig. S3C). Similarly, we find that RPLP2 increases in FXR1 overexpressing cells with AraC treatment (Fig. 3Cb, Table S3). Consistent with studies that demonstrated that P stalk proteins can interact with and activate GCN2 and cause eIF2α phosphorylation (*85–87, 94*), we tested FXR1 overexpression cells for GCN2 phosphorylation that marks its activation, and for downstream eIF2α phosphorylation. We find that FXR1 overexpressing cells—where RPLP0 increases as in AraC-treated cells (Fig. 3Ca)—show increased GCN2 phosphorylation and thus activation (Fig. 3Ea), and concordantly, increased eIF2α phosphorylation (Fig. 3Eb). Concurrently, we find that GCN2 associates more significantly with ribosomes (rRNA graph) and RPLP0 in FXR1 overexpressing cells (Fig. 3Fa), which may enable GCN2 activation and eIF2α phosphorylation. Consistently, this is reversed in FXR1 knockdown cells where phosphorylation of eIF2α decreases upon FXR1 depletion in THP1 cells (Fig. 3Fb), as well as on FXR1 depletion in other cell types (Fig. S3E). Importantly, GCN2 phosphorylation and activation as well as eIF2α phosphorylation in FXR1 overexpressing cells, is reduced upon depletion of RPLP0 (Fig. 3Fc), consistent with the need for RPLP0 in FXR1 overexpression conditions for GCN2 activation and eIF2α phosphorylation. These data suggest that ribosome changes in FXR1 overexpression cells, as in G0 and AraC treated cells where FXR1 increases, is associated with eIF2α kinase activation and eIF2α phosphorylation.

### FXR1 overexpressing cells, similar to G0 and AraC-treated cells, promote non-canonical translation

eIF2α phosphorylation inhibits canonical translation and reduces the stringency of selecting canonical AUG start sites that are in strong Kozak consensus sequence. This permits non-canonical translation, including use of mRNAs with complex or structured 5′ UTRs, or start sites embedded in poor Kozak consensus regions, or non-AUG start sites (*9, 95–103*). FXR1 increases in G0, and FXR1 overexpressing cells have multiple alterations of ribosome components, modifications, and complexes (Fig. 2, 3A-C). Increased snoRNA modification can promote non-canonical translation (*59*), altered ribosomes can promote specific mRNA and 5′UTR translation (*81*), and phosphorylation of eIF2α can inhibit canonical translation to permit non-canonical translation of specific peptides from non-canonical start sites (*99, 100, 102, 103*). Therefore, given that these ribosome and translation changes upon FXR1 increase in G0 and AraC-treated cells can re-direct translation, we tested luciferase reporters with a GUG start site that would be translated via non-canonical translation, over a canonical AUG start site reporter, and normalized to a Renilla reporter as co-transfection control.

In serum-starved G0 and AraC-surviving cells, we find that the translation of GUG reporter over that of the AUG reporter is enhanced, compared to the translation ratio in untreated, serum-grown proliferating cells (Fig. 3Ga), indicating that non-canonical translation is enabled in these G0 and AraC-treated conditions. Given that FXR1 is increased in these conditions, we asked if FXR1 overexpression affects non-canonical translation, as this translation could be due to other factors in G0 and AraC-treated cells or due the FXR1 increase in these conditions. We find that, as in G0 cells, in FXR1 overexpressing cells even without induction of G0 or treatment with AraC, the translation of GUG reporter over that of the AUG reporter is enhanced (Fig. 3Gb), compared to control vector expressing cells. These changes in luciferase expression are not seen at the RNA levels of the reporters (Fig. S3F). These data indicate that non-canonical translation is enabled by FXR1 increase, in FXR1 amplified cells, and in G0 and AraC-treated cells where FXR1 increases.

### FXR1 overexpression promotes translation of survival genes

The above data suggested that FXR1 overexpression in G0 and AraC-treated cells may promote non-canonical translation of specific mRNAs. We therefore performed polysome analysis of FXR1 overexpressing cells compared to vector control cells (Fig. 4Aa). The levels of subunits increase moderately (1.3 and 1.5 fold for 40S and 60S subunits, normalized to 80S) in FXR1a overexpressing cells, consistent with their decrease in FXR1 depleted cells in Fig. 1E. We analyzed the heavy polysomes by qPCR for mRNAs that are known to be increased in G0 THP1 cells but decreased upon FXR1 depletion from our datasets (Table S3, (*1, 28*)). Furthermore, these mRNAs have GC-rich 5′UTRs, as shown previously (*1, 104, 105*). As such GC-rich 5′UTRs are not preferred in canonical translation, these could be enabled by non-canonical translation conditions as in G0 and FXR1 overexpression cells where eIF2α is phosphorylated. These mRNAs include cell adhesion genes such as NCAM1, and anti-tumor immune regulators such as CD47 (*1, 106, 107*) that are implicated in chemosurvival, and increased in chemoresistant G0 cells and in AraC-treated cells that overexpress FXR1. Consistently, qPCR analysis of polysome fractions compared to monosomes that are normalized for input levels, reveal that these mRNAs are enriched on polysomes of FXR1 overexpressing cells compared to vector cells (Fig. 4B)—indicating their increased translation when FXR1 is increased, which could lead to chemosurvival of such FXR1 amplified cells.

**Fig. 4.**
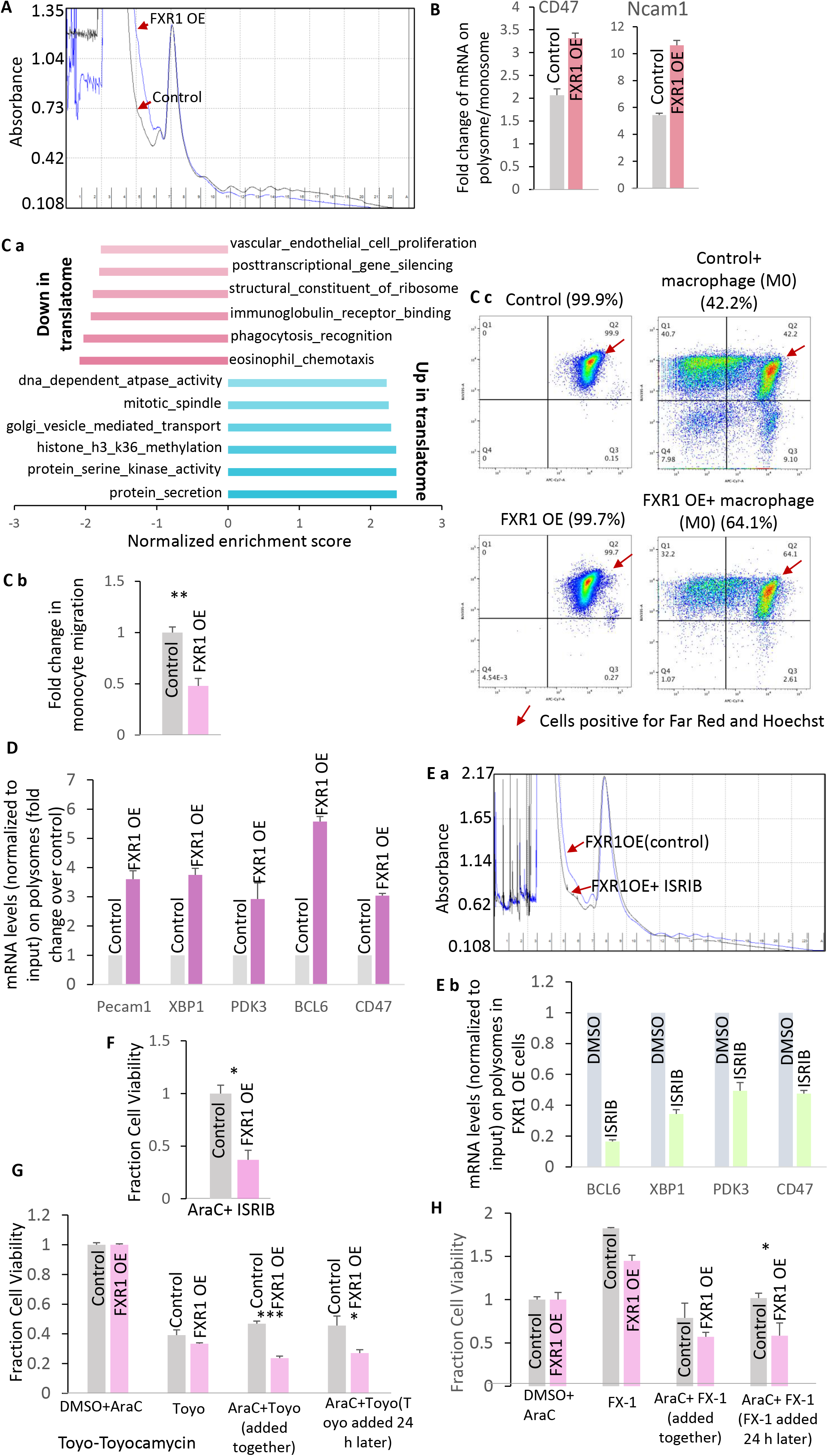
Chemosurvival in G0 and AraC-treated cells, and upon FXR1 overexpression, can be reduced by overriding eIF2α phosphorylation, and thereby decreasing non-canonical translation of pro-survival genes, or by targeting the translated pro-survival genes. **A.** Polysome analysis in FXR1 OE cells compared to control cells. **B.** qPCR analysis of polysomes of FXR1 OE cells compared to control cells, normalized for input levels, of pro- survival gene mRNAs that were previously found to increase in G0 and AraC-treated cell proteome in (*1, 28*). Ca. Graph showing gene ontology terms with highest and lowest Normalized Enrichment score (NES) as per GSEA analysis of the FXR1 OE translatome compared to control vector translatome from Table S4b-c. b. Graph of fold change in monocyte migration (image in Fig. S4Ca, stained with Far-red cell trace dye) in a trans-well assay with FXR1 OE cells or control vector cells in the bottom chamber. c. Flow cytometry data showing survival of control vector cells compared to FXR1 OE cells (stained with Far-red cell trace dye, as well as with Hoechst 33342 to detect live cells in quadrant 2 marked by an arrow) after co- culturing with macrophages (table of flow cytometry data of co-culture with monocytes, polarized, and resting macrophages in Fig. S4Cb). **D.** Polysome profiles in FXR1 OE compared to control cells. Shown qPCR analysis of selected mRNAs from FXR1 OE translatome compared to control vector translatome dataset in Table S4, normalized to inputs. E. a. Polysome analysis of FXR1 OE cells with or without ISRIB treatment b. followed by qPCR (graph) analysis of pro- survival gene mRNAs on polysomes. F. Chemosurvival of FXR1 OE cells, compared to control vector cells (gray bar), treated with AraC and ISRIB (co-treatment, restores canonical translation, suppressing non-canonical translation), **G.** with Toyocamycin, an XBP1 inhibitor (co-treatment with AraC, and post-AraC treatment), and H. with FX1, BCL6 inhibitor in FXR1 OE (post-AraC treatment) compared to control vector cells (gray bar), and to FXR1 overexpression and vector cells treated with AraC and buffer as controls. Data are average of 3 replicates +/-SEM. See also Fig. S4, Table S4.

We next asked globally which mRNAs are translated due to FXR1 overexpression, as also seen in G0 and in AraC-surviving cells where FXR1 increases, but without G0 or AraC treatment that could induce other effectors. Therefore, we conducted polysome profiling of vector and FXR1 overexpressing cells in untreated conditions without G0 or AraC treatment (Fig. 4A). The heavy polysomes (>2) were pooled and profiled by microarray compared to input samples to identify mRNAs that are promoted or repressed on polysomes by FXR1 overexpression (Table S4b-c).

We find that ∼10% (300 genes out of 2800) are genes that are commonly upregulated in the FXR1 overexpression cell translatome and in the increased proteome of AraC resistant cells (Table S4d). Apart from cell adhesion genes like PECAM1 (*108*), anti-tumor immune regulators such as CD47 (*106, 107*), and stress response genes like XBP1 (*109*) that we previously noted to be upregulated in G0 and AraC-treated cells (*1*) (Table S4a-e), mRNAs translated upon FXR1 overexpression also include pro-survival oncoproteins like BCL6 (*110–112*), signaling regulators that include the PI3K-AKT-mTOR pathway (*11, 113–117*), and metabolic enzymes like PDK3 (*118, 119*) that are known to promote tumor survival and progression (Fig. 4Ca, Table S4b-e).

These RNAs that are upregulated in FXR1 overexpression translatome that overlap with increase in AraC resistant cell proteome, have GC-rich 5′UTRs (Fig. S4A), as shown previously (*1, 104, 105*). Interestingly, several genes known to be translated by non-AUG start sites (*101*) are included among the upregulated translatome in FXR1 overexpression cells and have GC-rich 5′UTRs (Fig. S4B), which would be favored by non-canonical translation in G0, in AraC-treated cells, and in FXR1 overexpressing cells where eIF2α is phosphorylated. In addition, several classes of critical genes such as immune cell receptors as well as phagocytosis genes that include chemokines, are decreased in the translatome of FXR1 overexpressing cells compared to vector control cells (Fig. 4Ca, down in translatome genes, Table S4f), indicating specific gene expression to enable AML survival. These data suggest that specific gene categories are modulated by FXR1 increase to enable chemosurvival.

### FXR1 overexpressing cells show decreased immune cell migration, and increased survival with macrophages

Our data show altered translation of immune regulators in FXR1 overexpressing cells. Based on our data, these cells may subvert immune cells by decreasing translation of immune susceptibility genes such as immune receptors and phagocytosis related chemokine genes, while also promoting translation of immune evasion genes like CD47 that are known to inhibit anti-tumor activity by macrophages (*106, 107*) (Fig. 4Ca, up and down in translatome genes, Table S4d-f), which would lead to increased survival of such refractory AML cells.

Since FXR1 overexpressing cells show decreased translation of phagocytosis genes including chemokines (Fig. 4Ca, down in translatome genes, Table S4f), we first tested whether FXR1 overexpression alters migration of monocytes that can enable AML survival. We find that monocytes, co-cultured with control vector cells or FXR1 overexpressing cells in a trans-well assay, show 50% reduction in cell migration with FXR1 overexpressing cells compared to vector control cells (Fig. 4Cb, S4Ca). This is consistent with our data that show decreased translation of phagocytosis related chemokine genes in FXR1 overexpressing cells compared to vector control cells (Fig. 4Ca, down in translatome genes, Table S4f), as well as our previous data that showed altered immune gene regulators in G0 and AraC-treated cells (*1*). These data reveal that monocytes are precluded from the environment around FXR1 overexpressing cells through decreased chemokine translation, which could allow FXR1 overexpressing cells to evade immune phagocytosis, and associated anti-tumor immune activity.

Secondly, the immune evasion genes that we find increased in the translatome with FXR1 overexpression (Table S4d-e), are also increased seen in G0 and AraC-treated cells (*1*), and include genes such as SLAMF6 (*120, 121*), and CD47 (*106, 107*). These interact with immune cells to block anti-tumor immune activity (*122–124*). Therefore, we tested whether FXR1 overexpressing cells would show increased survival when co-cultured with macrophages that should be evaded by the presence of CD47, compared to vector control cells. Flow cytometry analysis revealed greater survival of FXR1 overexpressing cells (by 22%) compared to control vector cells, after co-culturing with macrophages (Fig. 4Cc, S4Cb). These data suggest that FXR1 overexpressing cells translate specific immune evasion regulators to survive anti-tumor immunity, indicating the role of FXR1 increase in G0 and AraC-treated cells. Together, these data suggest that the non-canonical translatome elicited in FXR1 amplified conditions, as in G0 and AraC-treated cells, decreases immune genes that would cause susceptibility to anti-tumor immune action such as chemokines, as well as promotes translation of anti-tumor immune evasion genes that inactivate anti-tumor immune activity.

### Chemosurvival of FXR1 overexpressing cells is reduced by drugs that override eIF2α phosphorylation, or block translated survival genes, BCL6 and XBP1

eIF2α phosphorylation is commonly observed in G0 cells and AraC-treated cells as previously noted (*1*) where FXR1 increases (Fig. 1A), and in FXR1 overexpressing cells (Fig. 3D-F). This increase in eIF2α phosphorylation is a potential vulnerability as such cells use non-canonical translation—elicited when eIF2α is phosphorylated—to express genes for chemosurvival. To test whether the increased eIF2α phosphorylation and non-canonical translation is needed for chemosurvival, we used the small molecule inhibitor ISRIB that is known to override eIF2α phosphorylation and restore canonical translation, which would thereby suppress non-canonical translation (*10, 125*). We first tested whether the translatome observed with FXR1 overexpression can be reversed by using ISRIB to restore canonical translation and block non-canonical translation. As shown in Fig. 4Ea, we performed polysome analysis in FXR1- ovexpression cells with or without ISRIB treatment followed by qPCR of the above identified mRNAs that are translationally increased in FXR1 overexpression cells. We find that ISRIB treatment reduced polysome association of mRNAs that are upregulated in FXR1 overexpressing cells (graph, Fig. 4Eb). These data support that FXR1 overexpression promotes non-canonical translation of mRNAs needed for chemosurvival. If so, then overriding this non-canonical translation should reduce the chemosurvival of FXR1 overexpressing cells. Consistently, we find that FXR1 overexpressing cells that are treated with both ISRIB and AraC have reduced chemosurvival compared to FXR1 overexpressing cells that are treated with buffer and AraC (Fig. 4F). To confirm that the translated genes are important for survival in FXR1 overexpression cells, chemosurvival was tested after inhibiting two known survival genes that are translationally upregulated in G0 and AraC treated cells, as well as in FXR1 overexpression cells, compared to vector control cells. We find that chemosurvival is decreased upon treatment with small molecule inhibitors of XBP1 and BCL6, two target genes that are translationally enhanced in G0, AraC-treated, and FXR1 overexpression cells: treatment of FXR1 overexpression cells with AraC and Toyocamycin (Fig. 4G, either together, or adding Toyocamycin post-AraC treatment), an inhibitor of XBP1, a pro-survival gene (*126–130*), or AraC and FX1 (Fig. 4H, adding FX1 post-AraC treatment), an inhibitor of BCL6 pro-survival oncoprotein (*110–112*), reduced chemosurvival compared to treatment with buffer and AraC, as well as compared to control vector cells. These data indicate that these translated genes are need for chemosurvival are a vulnerability that can be targeted in FXR1 overexpression conditions. Together, these data indicate that non-canonical translation mediated by FXR1 contributes to chemosurvival (Fig. S4D).

## Discussion

Cancer cells can enter a reversible arrest phase called quiescence or G0 that is resistant to harsh conditions including chemotherapy (*1–8*). Both G0 cells induced by serum-starvation, and post-chemo-treated surviving leukemic cells, are chemoresistant, and exhibit similar, specific gene expression at the post-transcriptional level (*1*). This indicated translation of a specific set of genes when cells become chemoresistant, upon surviving therapy, or upon entering the transient G0 state. How cells translate specific genes is not known and would involve translational changes and regulators. Interestingly, we find RNA- and ribosome-associated regulators are similarly altered in G0 and chemosurviving cells (*1*) (Fig. 1Aa-b), indicating that they may mediate this gene expression via mechanistic changes. FXR1 is an RNA-binding protein that was found associated with ribosomes (*15, 79*), and is a post-transcriptional regulator (*28*) that is amplified at its chromosome locus in several aggressive cancers, where it induces dysregulation of critical gene expression (*26*). Our data revealed that FXR1 (FXR1a isoform) is increased in G0 cells of specific cancers such as THP1 AML cells (*28*). Since G0 cells are chemoresistant (*1*), this indicates that FXR1 that is increased in G0 THP1 AML cells, could play a role in chemosurvival. In accord, we find that similar to G0 THP1 cells, AraC chemotherapy-treated THP1 and other AML cells, such as NOMO1, transiently upregulate FXR1 (Fig. 1Ac, S1A-B). These data suggested that FXR1 could be increased in some cancers treated with therapies like AraC, as it may be important for chemosurvival. Consistently, we find that overexpression of FXR1a isoform that is increased in G0 cells (*28*), promotes survival of THP1 cells treated with AraC by almost 2-fold, while knockdown of FXR1 reduces chemosurvival to less than 50% (Fig. 1B-C). These data suggest that FXR1 mediates chemosurvival in conditions like G0 and chemo-treated AML cells where it increases.

FXR1 was initially identified associated with the ribosome (*15*), and we had previously found that FXR1 associates with and promotes translation of specific mRNAs in G0 cells (*28*).

Therefore, the role of FXR1 in chemosurvival could involve translation regulation. To test this, we analyzed global translation in cells with or without FXR1 depletion by nascent translation labeling. We find that in FXR1 depletion cells, nascent translation is decreased by 50%, compared to control shRNA cells (Fig. 1D). Analysis of polysomes to investigate this effect revealed a sharp decrease of 40-60% of both ribosome subunits in FXR1 depleted cells, compared to control shRNA cells (Fig. 1E). These data suggest that FXR1 depletion affects ribosomes significantly. These reveal a role for the increased FXR1 in G0 and chemo-treated cells in ribosome regulation that may lead to the increased chemosurvival observed here.

FXR1 is known to be nucleolar, nuclear, and cytoplasmic, with roles in post-transcriptional regulation (*15–25*). Therefore, FXR1 can participate in ribosome biogenesis at multiple levels: through known associations with ribosomes and RPs (*15*), with RNA regulators, and RNAs that encode for ribosome regulators as observed previously (*28, 75*). Interestingly, we find that FXR1 depletion reduces all rRNAs including the precursor, and decreases many snoRNAs that increase in G0, as well as alters several RPs; conversely, FXR1a overexpression leads to their increase (Fig. 2A-C, 3A, 3C). Our data show that FXR1 regulates levels of snoRNAs that modify rRNAs (Table S1a-d), many of which are known to be involved in modifying critical sites that can affect translation (*60–66*). Consistent with the increase of snoRD63 and snoRA2A on FXR1a overexpression, we find increased modification at their rRNA target sites on 28S rRNA (Fig. 2Da-b); in accord, pseudouridylation is reduced along with alterations in other known rRNA modifications upon FXR1 depletion, as observed by RNA mass spectrometry (Fig. 2Dc, S2A, Table S1g). FXR1 depletion and overexpression also alters U3 and U8 that process rRNAs (*33, 37, 47-56*), which would lead to the observed alteration in mature rRNA levels (Fig. 2B).

To understand how FXR1 may enable such ribosome changes, we analyzed the FXR1 protein and RNA interactomes (Tables S2, S1e-f). We find that FXR1 not only associates with many snoRNAs (Table S1e-f), but also associates with snoRNA and ribosome biogenesis regulators, such as NOLC1 (*70–74*) (Fig. 2Ea), and other ribosome associated factors (Table S2, Fig. S2Ba). Consistent with the fact that both Pol I transcribed rRNAs, as well as Pol II transcribed snoRNAs and RPs are affected by FXR1, we find that FXR1 interacts with DDX21 (Fig. 2Eb), a regulator that promotes Pol I and Pol II transcription of ribosome genes (*68, 69*). In accord, we find that DDX21 overexpression but not of a control vector, partially rescues the reduced 45S rRNA precursor levels in FXR1 knockdown cells (Fig. 2Ec). This is consistent with regulation of ribosomes as DDX21 can also bind snoRNAs including U3 to promote their levels & functions, as well as turn on ribosome gene transcription via Pol I and Pol II (*68, 69, 131, 132*). Given that all 4 rRNAs are affected by FXR1 levels, and the 45S rRNA precursor of mature 18S, 28S, 5.8S rRNAs, from Pol I transcription is also affected along with the Pol III transcript, 5S rRNA, our data suggested that FXR1 may influence a common rRNA transcription factor for Pol I and Pol III or for all three RNA polymerases. Consistent with recent data showing FXR1 regulation of the levels of c-MYC (*75*) that regulates ribosome gene transcription from all three polymerases (*76, 77*), we find that c-MYC levels are increased with FXR1 overexpression (Fig. 2F).

Additionally, we find that FXR1 associates with the mRNA of POLR1D, a transcription factor associated with and needed for both Pol I and Pol III transcription; consistently, FXR1 depletion decreases while overexpression increases POLR1D (Fig. 2G, S2C). As POLR1D is needed for rRNA transcription from Pol I and Pol III, FXR1 regulation of POLR1D expression levels would impact all 4 rRNA levels (Fig. 2A). Together, these data reveal that FXR1 associates with or modulates multiple ribosome biogenesis regulators, which can lead to ribosome and translation changes upon FXR1 increase in G0 chemoresistant cells.

Collectively, these data suggest changes in the ribosome, upon FXR1 overexpression that is seen in G0 and AraC-treated cells (Fig. 2, 3A-B). Consistently, we find that ribosomes migrate differently, in two distinct complexes in G0 and AraC-chemoresistant cells compared to one in proliferating cells; importantly, this is also observed with FXR1 overexpression compared to control vector overexpression (Fig. 3C), indicating that FXR1 increase induces similar differences in ribosome complexes. Such ribosome changes can alter translation, which could enable the cells to adapt to chemotherapy and survive.

Such multiple changes on the ribosome can alter many downstream mechanisms to alter translation. One way that ribosome changes could alter translation, would be by activating stress signaling. Ribosomes not only function as translation machineries, but can also directly activate stress signaling pathways (*85–90*). This includes eIF2α kinases like GCN2 that is activated by stalled ribosomes (*133*) or by ribosomal P stalk proteins (*85–87, 94*), as well as the eIF2α kinase PKR that can be activated by snoRNAs (*91–93*). These kinases disable canonical translation via eIF2α phosphorylation, which permits non-canonical translation on specific mRNAs such as those with non-canonical start sites. In accord, we find that FXR1 overexpression leads to activation GCN2 and PKR eIF2α kinases (Fig. 3Da, 3Ea). Overexpression of FXR1 can increase specific snoRNAs that can bind and activate PKR, an eIF2α kinase that is a dsRNA binding protein (*91–93*), which then causes eIF2α phosphorylation. Consistently, we find that overexpression of FXR1-regulated snoRNAs, snoRD46 or snoRA2A, lead to increased eIF2α phosphorylation (Fig. 3Db) that can enable non-canonical translation. In accord, previous data had shown that overexpression of FBL that leads to altered snoRNA-mediated rRNA modification, induces non-canonical translation (*59*). Importantly, we find that P stalk ribosomal proteins are altered in levels in FXR1 overexpression cells as in G0 chemoresistant cells (Fig. 3Ca-b, Fig. S3C, Table S3a-b). Changes in the P stalk proteins can lead to GCN2 activation and eIF2α phosphorylation, via GCN2 interaction with RPLP0 (*85–87, 94*). Consistently, we find that overexpression of FXR1a promotes RPLP0 increase that is also seen in AraC-treated cells, and leads to increased GCN2 phosphorylation and activation and consequently eIF2α phosphorylation (Fig. 3Eb). P-stalk protein RPLP0 or uL10 associates with GCN2; consistently, in FXR1 overexpression cells where RPLP0 increases along with eIF2α phosphorylation, we find increased association of GCN2 with ribosomes and with RPLP0 (Fig. 3Fa). In accord, in FXR1 depleted cells, we find that eIF2α phosphorylation is decreased (Fig. 3Fb). Accordingly, depletion of RPLP0 in FXR1 overexpressing cells attenuated phosphorylation and activation of GCN2, and consistently decreased phosphorylation of eIF2α (Fig. 3Fc), indicating that RPLP0 is needed for the eIF2α phosphorylation observed in FXR1 overexpression conditions. Together, these data suggest that increased FXR1 in G0 chemosurviving cells alters the ribosome, which then can induce stress signals to inhibit canonical translation and permit non-canonical translation.

Such stress signaling by the ribosome has been observed with colliding ribosomes that triggers either GCN2-mediated eIF2α phosphorylation that leads to survival, or at increased levels induces stress JNK/p38 MAPK signaling pathway to trigger apoptosis, and has also been observed with c-GAS signaling in other conditions, as well as with ribotoxic stress agents, leading to post-transcriptional gene expression changes (*88, 134–139*). With eIF2α phosphorylation, stringent use of conventional Kozak start sites for initiator tRNA recruitment (*140*) is reduced and can lead to non-canonical start site translation (*97, 99, 141-143*). This could permit translation of specific mRNAs such as those with complex 5′UTRs, unconventional non-AUG start sites, or AUGs in poor Kozak start sites that would normally be poorly translated by canonical translation (*99*). In accord, we find that translation of a reporter with a non-canonical GUG start site over that of a reporter with a conventional AUG start site, is promoted in FXR1 overexpressing cells, and in G0 and chemo-treated cells where FXR1 increases and elicits ribosome and phospho-eIF2α changes, compared to untreated serum-grown proliferating control vector cells (Fig. 3Ga-b, S3E). Consistently, we find that many of the genes upregulated in G0 and AraC-treated cells, as well as in FXR1 overexpressing cells (Fig. 4A-C, Table S4a-e) have GC-rich 5′UTRs and motifs (Fig. S4A). Additionally, several reported non-AUG start site genes (*101*) are translationally upregulated in these conditions of FXR1 overexpression (Table S4, Fig S4B). Other features that are unique to these transcripts such as previously identified 3′UTR elements (*1, 28, 144*), as well as other ribosomal and translational changes, may also contribute to the specific translation observed in these non-canonical translation conditions. Together, our data suggest that FXR1 increase in G0 and chemo-treated cells enables non-canonical translation, through a signaling role elicited by ribosome changes that lead to eIF2α phosphorylation and ISR.

Such inhibition of canonical translation by eIF2α phosphorylation can lead to expression of specific mRNAs that are critical under stress conditions, and are usually translationally restricted, either due to unconventional start sites, or GC-rich 5′UTRs that are poorly translated by the canonical mechanism (*96, 99, 100, 142, 145, 146*). We find that genes that were previously identified as translated in G0 chemoresistant cells (*1*) show enhanced polysome association in FXR1 overexpressing cells (Fig. 4A-C, Table S4b-e). These genes are known targets downstream of ISR such as XBP1 (*147, 148*), or have high negative ΔG with a GC-rich motif in their 5′UTRs in genes such as PECAM1 (*108*), CD47 (*1*), as well as NCAM1 that is known to have a structured 5′UTR (*104*) (Fig. S4A); these indicate that non-canonical translation conditions could enable their increased translation, as previously observed with such 5′UTRs (*105*). Consistently, treatment with ISRIB, an ISR small molecule inhibitor that overrides eIF2α phosphorylation to restore canonical translation (*10, 125*), prevents this non-canonical translation of the identified genes (Fig. 4D-E). Other ribosome changes elicited by FXR1 increase may also contribute to altering translation, and effectors other than FXR1 and previously identified UTR binding and translation factors in G0 and AraC-treated cells (*1, 28, 144*) may also contribute to the specialized translation in these conditions. Thus, these data suggest that FXR1 increase in G0 and AraC-treated cells causes ribosomal changes that can reduce canonical translation and enable non-canonical translation of specific genes, by triggering eIF2α phosphorylation.

Importantly, our data reveal that pro-survival genes with critical roles in tumor survival and progression, such as stress response genes like XBP1 (*126–130*), cell adhesion genes like PECAM1 (*108*) and NCAM1 (*149*), immune genes like CD47 that promote AML survival (*106, 107*), as well as oncoproteins like BCL6 (*110–112*) (Table S4), are translated via these ribosome changes that are triggered by FXR1 increase in G0 chemosurviving cells. Other genes are downregulated in the FXR1 overexpressing cell translatome compared to control vector cells, including those that would increase susceptibility to anti-tumor immune action, including immune receptor genes and phagocytosis chemokine genes (Fig. 4Ca, down in translatome genes, Table S4f). Consistently, we find that monocyte migration is reduced by 50% in trans- well assays with FXR1 overexpressing cells compared to vector control cells (Fig. 4Cb, S4Ca), which could allow FXR1 overexpressing cells to evade immune phagocytosis, and associated anti-tumor immune activity. Concurrently, the increased translatome in FXR1 overexpressing cells include anti-tumor immune evasion genes such as CD47 (Fig. 4A, Table S4b-e) that can inactivate the anti-tumor immune response of macrophages to promote AML survival (*106, 107*). In accord, we find that FXR1 overexpressing cells show increased survival compared to control vector cells when co-cultured with macrophages (Fig. 4Cc, S4Cb), implicating immune survival of cells where FXR1 increases, such as in G0 and chemo-treated AML cells. Together, these data suggest that the non-canonical translatome elicited in FXR1 amplified conditions, as in G0 and AraC-treated cells, subverts anti-tumor immunity, by decreasing immune genes that would cause susceptibility to anti-tumor immune action such as chemokines, as well as by promoting translation of anti-tumor immune evasion genes that inactivate anti-tumor immune activity.

Together, our data suggested that the enhanced chemosurvival in FXR1 overexpressing cells—as in G0 and chemoresistant cells, where FXR1 is increased and causes ribosome changes and eIF2α phosphorylation—could be reversed by targeting the eIF2α phosphorylation-induced non-canonical translation, or such downstream translated genes. Consistently, we find that inhibition of this non-canonical translation with ISRIB (Fig. 4F), or inhibition of the downstream target genes such as that of XBP1 (*127, 129, 130*) or of BCL6 (*110–112*) with pharmacological inhibitors (Fig. 4G-H), reduces the chemosurvival observed in FXR1 overexpression cells, compared to control vector cells or to FXR1-overexpression cells treated with chemotherapy alone. These changes could also potentially support the translational changes in aggressive cancers where FXR1 increases (*26*), and consistently, can be targeted therapeutically. Together, our data reveal that FXR1 is a critical translation regulator that is amplified in G0 and chemosurviving leukemic cells, which induces ribosome changes that trigger stress signals to enable non-canonical translation of pro-survival genes to promote tumor persistence (Fig. S4D).

## Acknowledgements

Raw datasets will be made available on the public repository, GEO, and all materials will be made available publicly on publication and on request. The study is funded by GM134944, & CA220103 grants from NIH and a Kurt Isselbacher grant from the RICBAC foundation to SV.

## Author Contributions

CD conducted the research; SST, SIAB, HN, BB, KQW, OLT, SL, SK, & MG contributed to the data; MB, JK, & WH conducted proteomics; SV supervised the project & wrote the manuscript. We thank S. Kollu for support on the data, D. Scadden and D. Sykes for cell lines, HSCI-CRM Flow Cytometry Core facility at MGH for flow cytometry analysis, Partners Healthcare Center for Personalized Genetic Medicine & BUMC facilities for microarray data, and ArrayStar for LC-MS modification data.

## Data and Materials Availability

Raw datasets will be submitted to GEO public repository at final submission and along with all materials will be available publicly as well as on request to the authors.

## List of supplementary Materials

Materials and Methods

Table S1-S5

Fig S1-S4

References

