## Supplemental Materials for "Ribosome changes elicit non-canonical translation for chemosurvival in G0 leukemic cells"

**List of supplementary Materials:**

Materials and Methods
Table S1-S5
Fig S1-S4
References

**S1A**

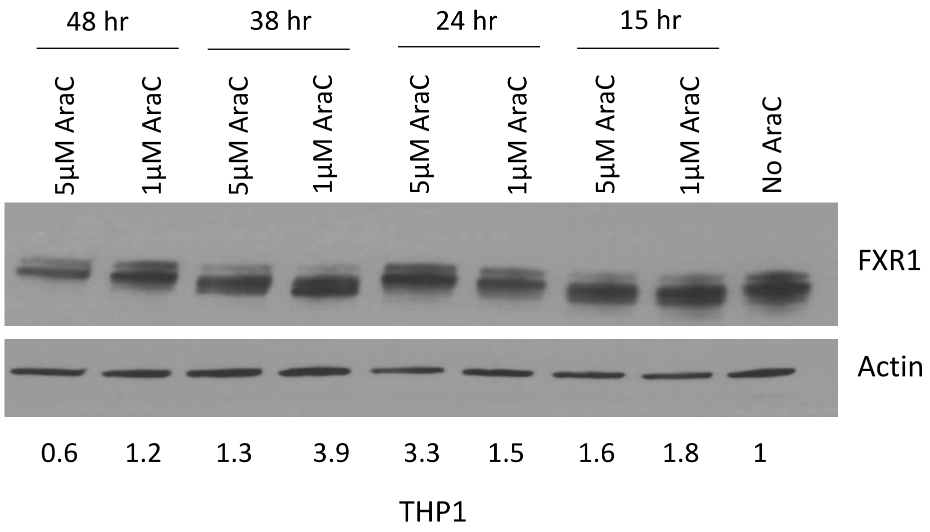
48 hr
38 hr
24 hr
15 hr
5μM AraC
1μM AraC
5μM AraC
1μM AraC
5μM AraC
1μM AraC
5μM AraC
1μM AraC
No AraC
FXR1
Actin
0.6
1.2
1.3
3.9
3.3
1.5
1.6
1.8
1
THP1

**S1B**

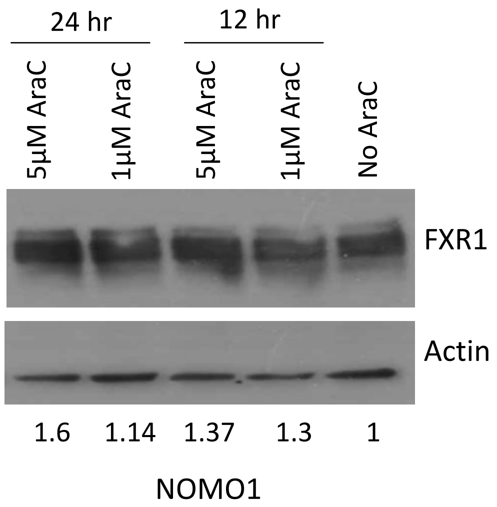
24 hr
12 hr
5μM AraC
1μM AraC
5μM AraC
1μM AraC
No AraC
FXR1
Actin
1.6
1.14
1.37
1.3
1
NOMO1

**S2A**

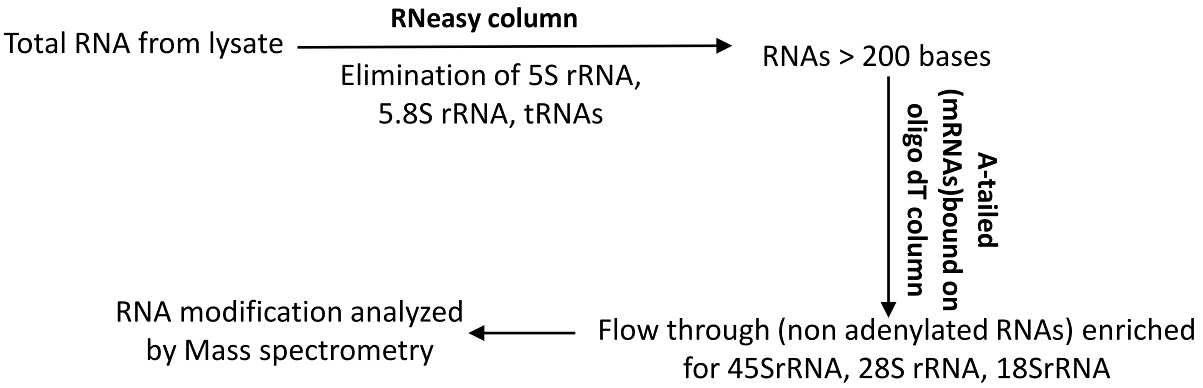


| RNA Mass Spec modification | Modified/Unmodified (RNA from FXR1 KD wrt Control) |
|---|---|
| Adenosine (Unmodified) | |
| 2'-O-methyladenosine(Modified) | 1.108472671 |
| Cytidine (Unmodified) | |
| 2'-O-methylcytidine(Modified) | 0.97898797 |
| Guanosine (Unmodified) | |
| 2'-O-methylguanosine(Modified) | 1.867703004 |
| Uridine (Unmodified) | |
| 2'-O-methyluridine(Modified) | 1.321412023 |
| Uridine (Unmodified) | |
| Pseudouridine(Modified) | 0.706883806 |
| Cytidine (Unmodified) | |
| N4-acetylcytidine(Modified) | 2.288172166 |

**S2B** **a**

| GeneName | FXR1 Antibody/IgG |
|---|---|
| EIF2B5 | 2.545751014 |
| EEF1A1 | 2.098479339 |
| RPL37 | 2.049285418 |
| RPL19 | 1.947400807 |
| RPS28 | 1.915770681 |
| NOLC1 | 1.900645092 |
| RPL36A | 1.8336335 |
| RPS6KA3 | 1.794537553 |
| RPS3A | 1.787639249 |
| RPS17 | 1.752414395 |
| NAT10 | 1.734520121 |
| RPS14 | 1.680766661 |
| RPL18A | 1.646411455 |
| RPL10A | 1.566336262 |

**b** **FXR1 IP from cytoplasmic fraction**

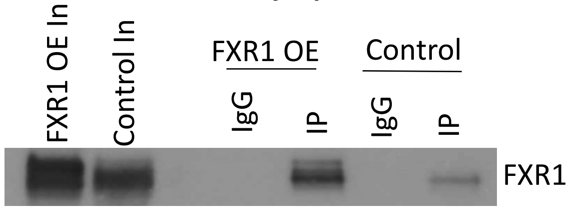


**Binding of FXR1 to rRNAs**

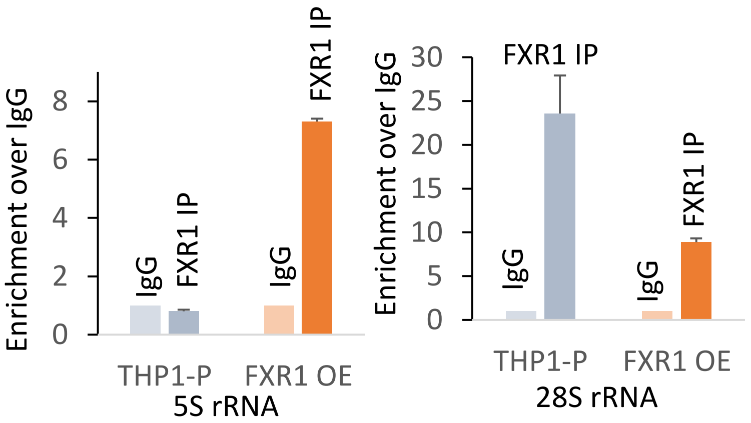


**S2C**

| Condition | Gene Name | FXR1KD/ Control |
|---|---|---|
| S+ | PolR1D iso 2 | 0.703402242 |
| 1DG0 | PolR1D iso2 | 0.732959199 |

**S3A**

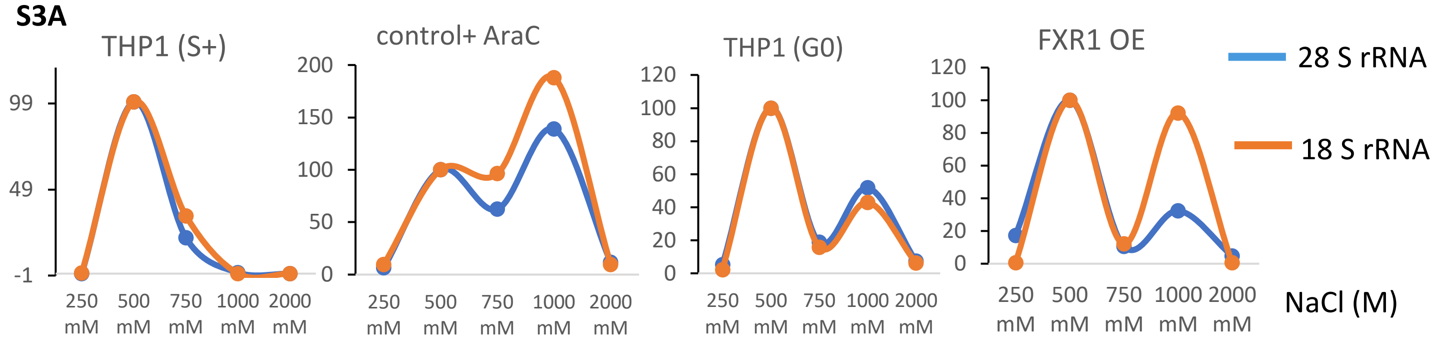


**S3B a**

**Ribosome IP from cytoplasmic extracts**

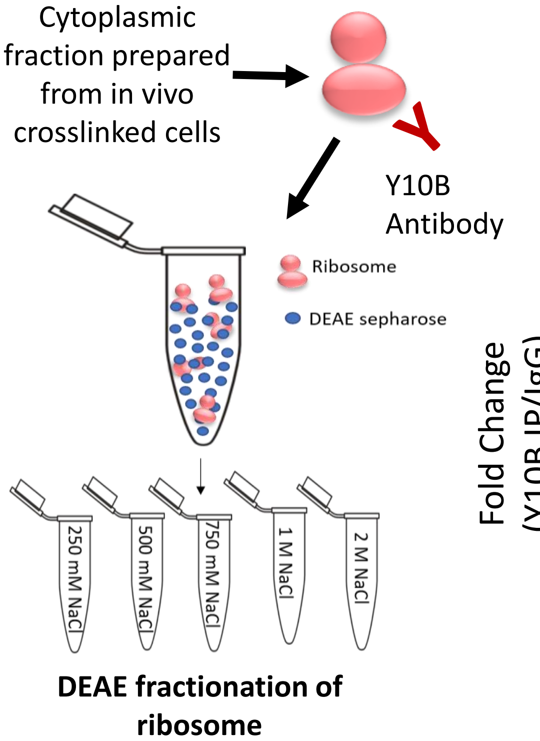


**S3B b**

**Y10B IP from cytoplasmic fraction**

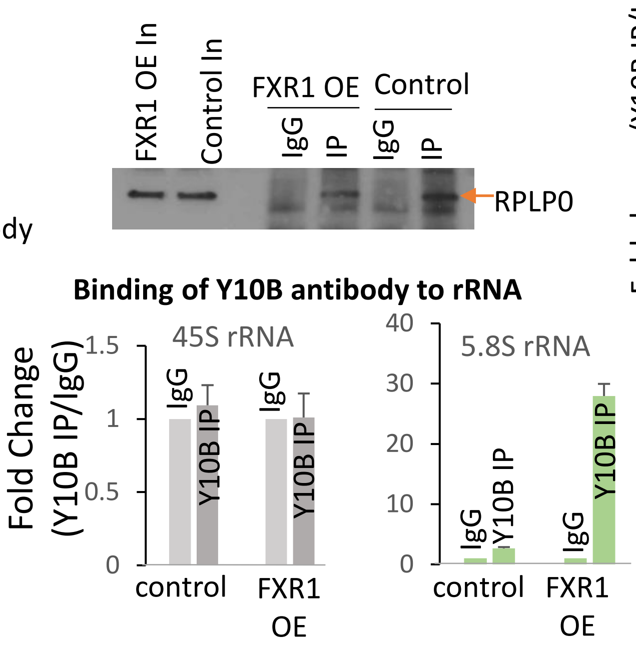


**S3B c**

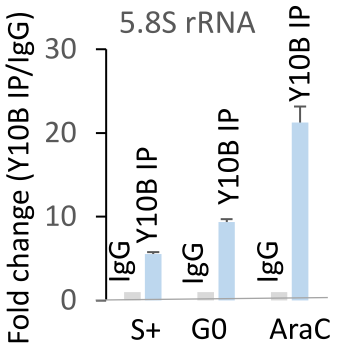


**S3C**

| Protein | mRNA level microarray (FXR1 KD/Control) | mRNA level in translatome (FXR1 KD/Control) | Protein level (FXR1 KD/Control) |
|---|---|---|---|
| RPLP0 | -1.32 | -1.32 | 0.719335688 |

**S3D**

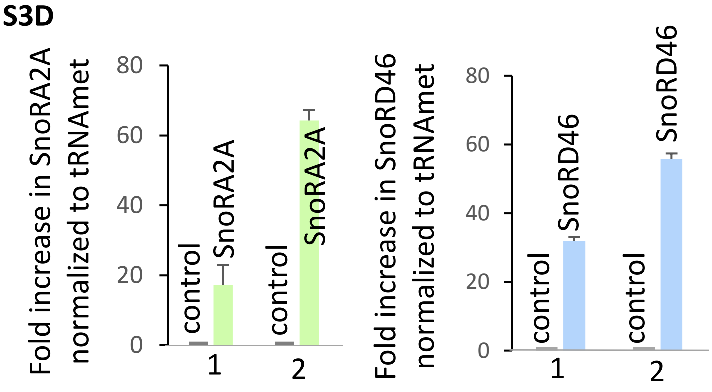


1 and 2 represent biological duplicates

**S3E** **FXR1 KD in BT549**

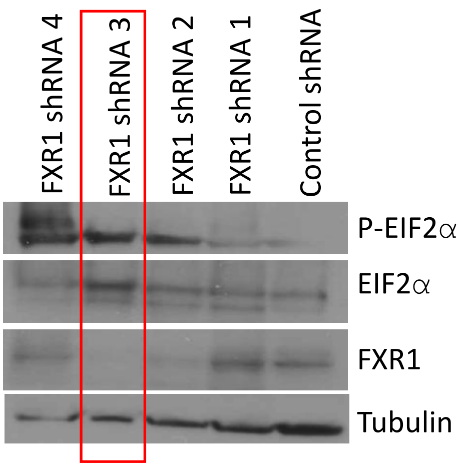


**S3F**

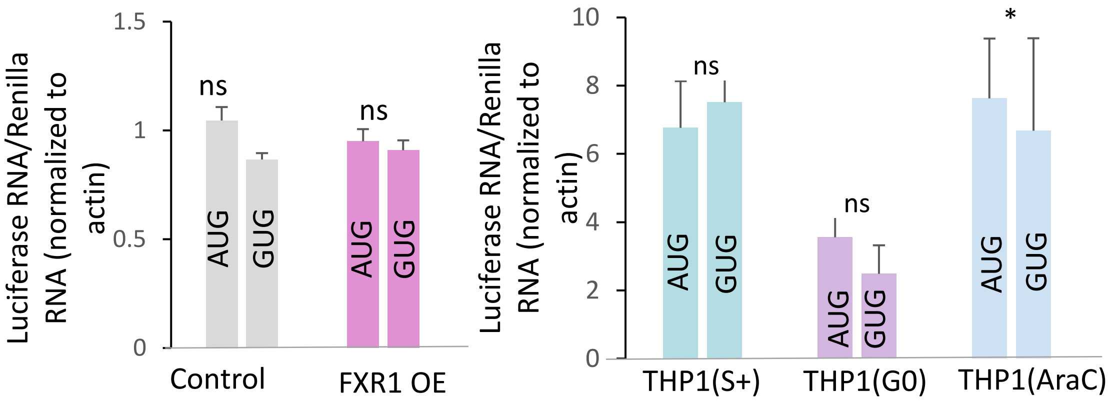

S4A a

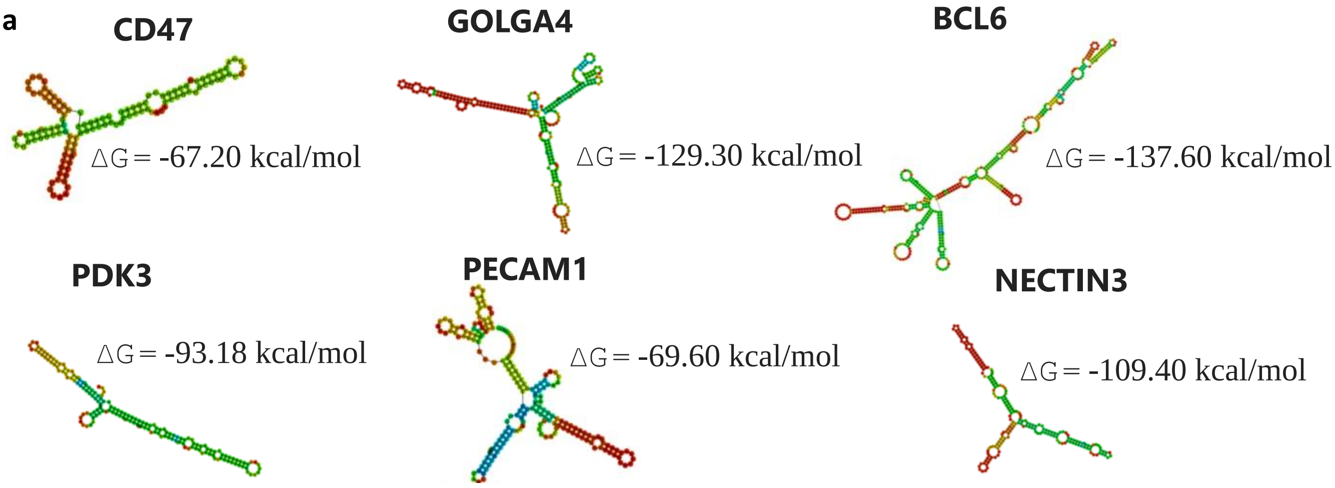


b

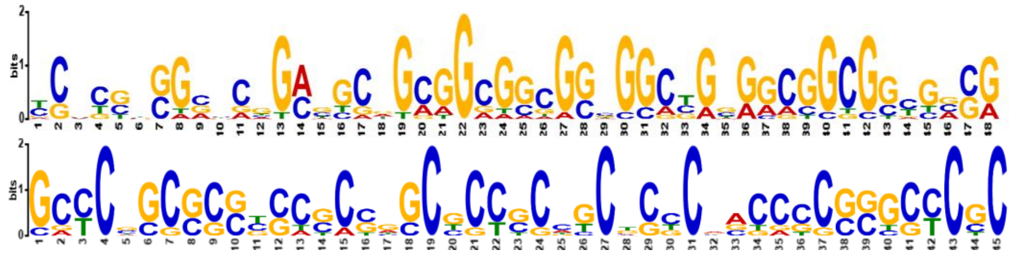

S4B a

| Gene name | Fold change on polysome normalized to Input (FXR1 OE/Control) | Translation start site |
|---|---|---|
| TEAD3 | 1.6 | AUA |
| CITED1 | 1.6 | CUG |
| STARD10 | 1.5 | GUG |
| GTF3A | 1.4 | CUG |
| VEGFA | 1.4 | CUG |
| EIF4G2 | 1.3 | GUG |
| NPW | 1.3 | CUG |
| NFKBID | 1.3 | CUG |
| WDR26 | 1.3 | ACG |
| RNF187 | 1.2 | CUG |
| EIF4G3 | 1.2 | AUC |
| YPEL2 | 1.2 | ACG |
| TLE3 | 1.2 | CUG |
| ZFP62 | 1.2 | GUG |
| EPHX3 | 1.2 | ACG |

b

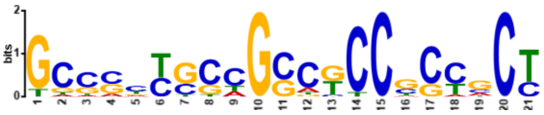

S4C a

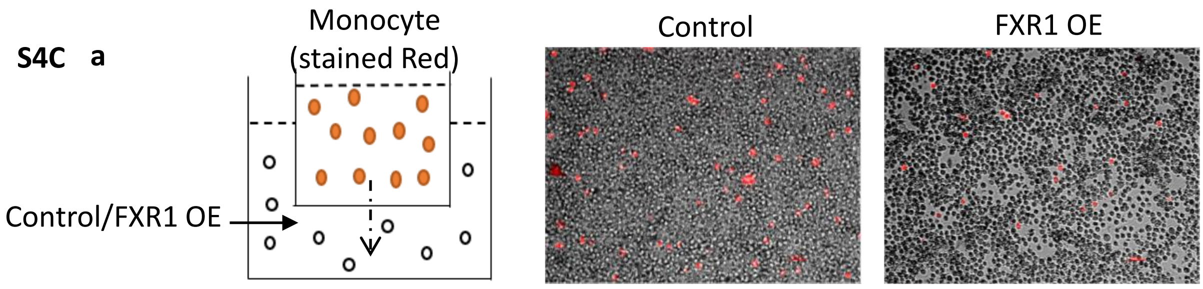


b

| Test cells | Cocultured with | % survival of Test cells |
|---|---|---|
| Control | - | 99.9% |
| FXR1 OE | - | 99.7% |
| Control | Monocyte polarized to M0 | 42.2% |
| FXR1 OE | Monocyte polarized to M0 | 64.1% |
| Control | Monocyte polarized to M1 | 74.9% |
| FXR1 OE | Monocyte polarized to M1 | 79.5% |
| Control | Monocyte polarized to M2 | 46.0% |
| FXR1 OE | Monocyte polarized to M2 | 65.4% |
| Control | Monocyte | 35.4% |
| FXR1 OE | Monocyte | 44.6% |

**S4D**

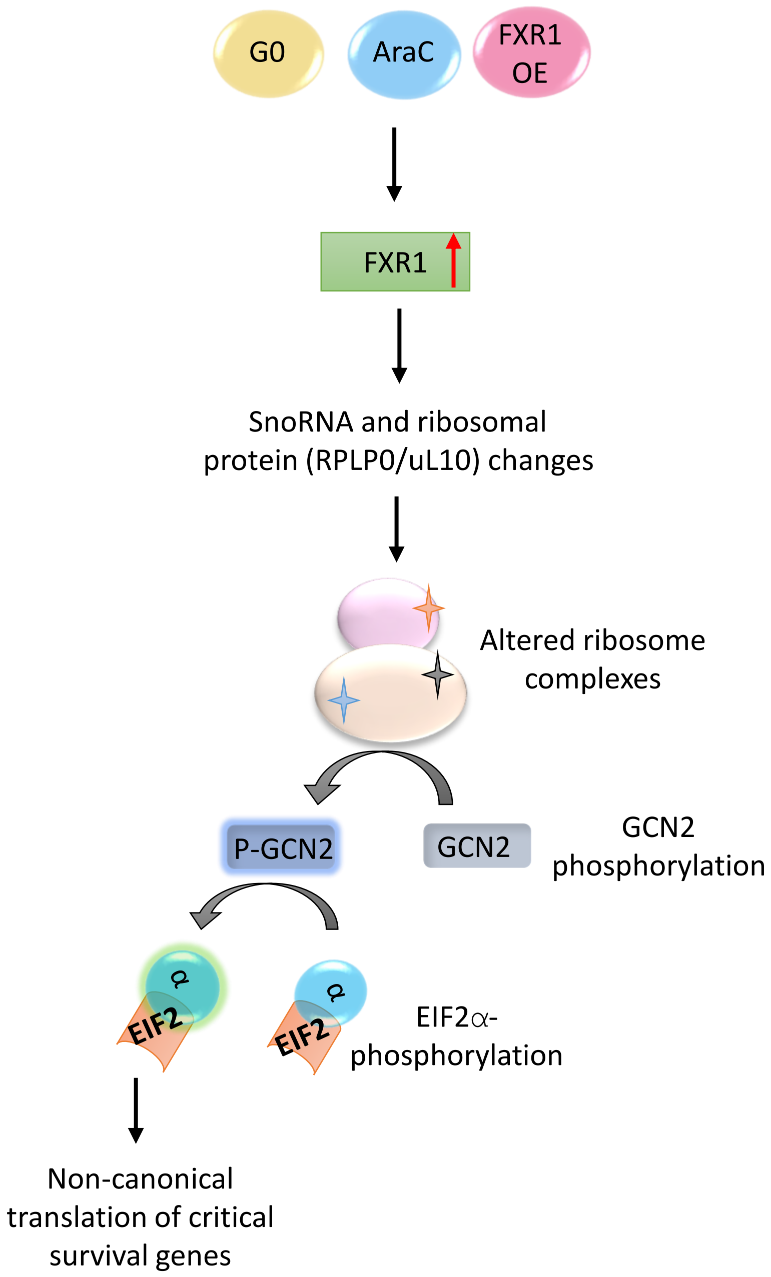
G0
AraC
FXR1 OE
FXR1
SnoRNA and ribosomal protein (RPLP0/uL10) changes
Altered ribosome complexes
P-GCN2
GCN2
GCN2 phosphorylation
α
EIF2
α
EIF2
EIF2α-phosphorylation
Non-canonical translation of critical survival genes

**Materials and Methods:**

**Cell Culture**

THP1 cells were cultured in Dulbecco's modified Eagle medium RPMI1460 media supplemented with 10% fetal bovine serum (FBS), 2 mM L-Glutamine, 100 µg/mL streptomycin and 100 U/ml penicillin at 37°C in 5% $CO_2$. SS or G0 THP1 cells were prepared by washing with PBS followed by serum-starvation at a density of $2 \times 10^5$ cells/mL and AraCS cells, by treatment with indicated concentrations of AraC for indicated periods of time. THP1 (TIB-202) and monocytes (CRL9855) were obtained from ATCC. NOMO1 and MOLM13 were obtained from ATCC and from DSMZ by the Scadden group (*1*). Cell lines were tested for Mycoplasma (Promega) and authenticated by the ATCC Cell Authentication Testing Service (*1*).

**Plasmids**

TRIPZ and GIPZ plasmids expressing shRNAs against human FXR1, RPLP0, and control vector expressing miR30a primiR sequences (RHS4750), were obtained from Open Biosystems-Dharmacon (shRNA target sequences are in Table S5). Stable cell lines were constructed as described previously (*2, 3*). The stable cells expressing shRNA against FXR1 were induced with 1 µg/mL doxycycline for three days (once each day) to knockdown FXR1. Control cells were treated similarly. THP1 FXR1 OE cell lines were created by transducing cells with pHAGE retroviral vector containing FXR1a for constitutive over expression of FXR1a (*3-6*). Renilla was obtained and used previously (*4*). DDX21 plasmid (*7*) and GUG and AUG Luciferase reporters (*8*) were obtained from Addgene.

**Polysome profiling and microarray**

30 x 10^6 cells were grown for each sample and harvested on ice. In case of the FXR1 KD and its respective control, cells were treated with 100 ug/ml of cycloheximide (Sigma 66-81-9) for 5 minutes prior to harvesting. In case of FXR1 OE cand its respective control untreated or treated with ISRIB, cycloheximide was not used. Sucrose was dissolved in lysis buffer containing 100 mM KCl, 5 mM $MgCl_2$, 2 mM DTT and 10 mM Tris-HCl (pH 7.4). Sucrose gradients from 10% to 55% were prepared in ultracentrifuge tubes (Beckman) as previously described(*2, 4, 9, 10*). Harvested cells were rinsed with ice-cold PBS and resuspended in lysis buffer with 1% Triton X-100 and 40 U/mL murine (New England Biolabs) for 20 minutes with intermittent tapping on ice. After centrifugation of cell lysates at 12,000 x g for 20 minutes, supernatants were loaded onto sucrose gradients followed by ultracentrifugation (Beckman Coulter Optima L90) at 32,500 × rpm at 4 °C for 80 min in the SW40 rotor. Samples were separated by density gradient fractionation system (Biocomp Piston Gradient Fractionation). 100 ug/ml of cycloheximide was added to lysis buffer and sucrose solutions in case of cycloheximide treatment of cells prior to harvesting. RNAs were purified from heavy polysome fractions and whole cell lysates. The synthesized cDNA probes from WT Expression Kit (Ambion) were hybridized to Gene Chip Human Transcriptome Array 2.0 (Affymetrix) and analyzed by the Partners Healthcare Center for Personalized Genetic Medicine Microarray and BUMC facilities. Gene ontology analysis for differentially expressed translatome or proteome was conducted by DAVID 6.7 tools (*11*) (*12*). Molecular signatures enriched in FXR1 OE and Control cells were identified by Gene Set Enrichment Analysis (GSEA) (*13*).

**Western blot analysis**

Cells were collected and resuspended in lysis buffer containing 40 mM Tris-HCl (pH 7.4), 6 mM $MgCl_2$, 150 mM NaCl, 0.1% NP-40, 1 mM DTT, 20 mM, 17.5 mM ß-glycerophosphate, 5 mM NaF and protease inhibitors. Cell lysates were heated at 95°C with 200 mM DTT and 1X SDS loading dye for 10 min. Samples were loaded onto 4%-20% gradient SDS-PAGE (Bio-Rad) or 16% SDS-PAGE (Invitrogen), transferred to PVDF membranes and processed for immunoblotting. Antibodies against FXR1 (used for Western) (05-1529), Actin (MAB1501), Tubulin (05-829) were from Millipore; NOLC1 (11815-1-AP), DDX21(10528-1-AP), PolR1D (12254-1-AP), RPLP0 (11290-2-AP), RPLP2 (16805-1AP), RPL29 (15799-1-AP), RPL11 (16277-1AP), FXR1 (13194-1-AP) (used for Immunoprecipitation) were from Protein Tech; PKR (3072S), P-PKR (2611S), EIF2α (9722S), P-EIF2α (3597S), GCN2 (3302S) (used for immunoprecipitation) were from Cell Signaling; c-Myc (ab32072), P-GCN2 (ab75836), GCN2 (ab134053) (used for Western blot) were from Abcam.

**Mass Spectrometry**

Multiplex quantitative proteomics analysis (tandem-mass-tag (TMT) spectrometry) was conducted, as previously (*14*), from THP1 leukemic cells and cell lines created that were treated as described.

**Quantitative RT-PCR**

Total RNA was extracted using proteinase K buffer and TRIzol (Invitrogen) as performed previously (*3*). The cDNA was synthesized from 1 µg of RNA using M-MuLV Reverse Transcriptase (NEB) and random hexamer primer (Promega). qPCRs were run on LightCycler®

480 Instrument II (Roche) using 2 X SYBR green mix (Bio-rad). All primers used are listed in Table S5.

**Mass Spectrometry for RNA modification analysis**

Total RNA was isolated using Trizol as per manufacturer's instructions. Isolated RNAs were cleaned using RNeasy kit (Qiagen). Poly(A) containing RNAs were separated from the RNA pool using a Poly(A) mRNA isolation system IV (PolyATract, Promega). The remaining non-polyadenylated RNA was sent for nucleoside digestion and LC MS/MS analysis to Arraystar Inc (https://www.arraystar.com/lc-ms-based-rna-modification-analysis-service-selected/isolated-rna/).

**Nascent translation level analysis**

Global translation was measured by metabolic labeling for a short period followed by PAGE and scintillation analysis (*2*). THP1 stable cell lines were grown in normal RPMI medium to prevent additional cellular stress from methionine-free medium. 100 µCi of $^{35}$S-methionine was added to 10 mL of cells. After incubation at 37 °C for 45 min, cells were washed once with PBS and lysed in buffer (40 mM Tris-HCl (pH 7.4), 6 mM $MgCl_2$, 150 mM NaCl, 0.1% NP-40, 1 mM DTT and protease inhibitors)**.** The lysate was first separated by electrophoresis on an SDS-PAGE gel, then transferred to a PVDF membrane by Western Blotting that were exposed to a phosphoimager (GE Healthcare) and quantified by ImageJ. The lysates were also measured by scintillation counter (shown).

**Low dNTP RT-qPCR Assay**

For analysis of 2'*O*- methylation, RNA was prepared from indicated cells. Reverse transcription (as described above in qPCR) was performed with primers in the reverse orientation from the modification site on ribosomal RNA. Two different dNTP concentrations were used for each primer, low (0.025 mM) and high (2.5 mM). qPCR was done with the resulting cDNA using primers at forward and reverse orientation to the modified sites. Fold change in modification was calculated based on difference of $C_t$ values in the low and high dNTP conditions (*15*). For analyzing pseudouridylation, first the RNA was treated with CMC metho-*p*-toluene sulfonate (1-cyclohexyl-(2-morpholinoethyl) carbodiimide metho-*p*-toluene sulfonate) under alkaline conditions (*15-18*). This was followed by the process for low dNTP RT-qPCR, as described above for 2'-*O*- methylation. Primers are listed in Table S5.

**In vivo crosslinking and Immunoprecipitation**

In vivo crosslinking using 0.3% formaldehyde, and nuclear-cytoplasmic separation were done as described earlier (*4, 5*). Briefly, around 10-15 x 10^6 cells were harvested on ice followed by washing with cold PBS. Cell pellets were resuspended in hypotonic buffer (10 mM Tris (pH=8), 1.5 mM $MgCl_2$, 10 mM KCl). Cells were lysed using a syringe with 25G*5/8 precision glide needle. Cytoplasmic fraction was collected by centrifuging at 2000 rpm for 10 mins. The remaining pellet was resuspended in equal volumes of buffer (20 mM Tris (pH=8), 25% glycerol, 1.5 mM $MgCl_2$, 0.2 mM EDTA, 20 mM KCl) and buffer (20 mM Tris(pH=8), 25% glycerol, 1.5 mM $MgCl_2$, 0.2 mM EDTA, 1.2 mM KCl), incubated on cyclomixer for 45 min, sonicated 6 times, 30 secs each, centrifuged for 10 min at 2000 rpm. The supernatant was mixed with the cytoplasmic fraction. Where mentioned, only cytoplasmic fraction was used. The lysates were pre-cleared of non-specific binders by incubating with Protein G (Santa Cruz Biotechnology) beads and IgG. The pre-cleared lysates were incubated with antibodies and IgG as controls overnight in buffer (40 mM HEPES, 100 mM NaCl, 6 mM MgCl2, 0.025% NP-40, 1 mM DTT, 10% glycerol, 1mM PMSF). Lysates were then incubated with equilibrated and blocked Protein G beads for 2 hr. Beads were then pelleted and washed 4 times with RIPA buffer. Beads were then either used to analyze proteins or RNA after using heat to break Schiff's linkages from formaldehyde followed by proteinase K digestion buffer treatment for the fractions for RNA analysis, and RNase treatment for the fraction for protein analysis as described earlier (*4, 5*).

**DEAE (diethylaminoethanol) fractionation assay**

DEAE fractionation was performed with in vivo formaldehyde crosslinked extracts or Y10B antibody immunoprecipitates as developed previously (*4*). Cell lysates or fractions post Y10B immunoprecipitation, were incubated with equilibrated DEAE beads for 2 hr in buffer (40 mM HEPES, 6 mM $MgCl_2$, 2 mM DTT, 10% glycerol, 150 mM NaCl). After collection of the flowthrough, beads were incubated with wash buffer of increasing salt concentrations (40 mM HEPES, 6 mM $MgCl_2$, 2 mM DTT, 10% glycerol, with NaCl ranging from 250 mM-500 mM-750 mM-1000 mM-2000 mM). RNA was isolated from each salt fraction. Amount of ribosomal RNA in each fraction was analyzed by qRT-PCR, normalized to input levels.

**Luciferase Assay**

Plasmids containing firefly luciferase reporters downstream of AUG/GUG start sites (*8*) were co-nucleofected using Lonza Nucleofector (*3*) with Renilla luciferase in FXR1 OE, control vector, and THP1 cells. Nucleofected cells were then grown under conditions of S+, G0, and 5 μM AraC. Cells were harvested, washed, and lysed in 1X passive lysis buffer (Promega). Luciferase activity in the lysates was analyzed using Luciferase Assay System (Promega) as per manufacturer's instructions and as conducted previously (*4*).

**Co-culture of leukemic cells with immune cells followed by flow cytometry**

CRL9855 monocytes were tested as monocytes or polarized to M0, M1, M2 macrophages with 185 ng/ml of PMA (phorbol 12-myristate 13-acetate) for 24 hr and continued to derive M0 macrophages. M0 macrophages were then treated with 20 ng/ml of interleukin 4 and 20 ng/ml of interleukin 13 for 48 hr for polarization to M2, and with 20 ng/ml of Interferon Y and 20 ng/ml of TNFα for 48 hr for polarization to M1 (*19, 20*). Control and FXR1 OE cells were stained with Far-Red Cell Trace dye (Thermo Fisher Scientific) as per manufacturer's instructions. Macrophages were co-cultured with stained control and FXR1 OE cells for 12 hr at a ratio of 1:2. Co-culture cells were collected after trypsinization and washed with PBS. Non-co-cultured control/FXR1 OE cells stained with Far Red Cell Trace were used as controls. Harvested cells were thoroughly washed and stained with Hoechst 33342 (Thermo Fisher Scientific) as per manufacturer's instructions. Samples were filtered through a nylon mesh filter and analyzed for the population stained with both dyes by Flow cytometry (*21*). Control cells without the Far-Red Cell trace but stained with Hoechst were used as baseline and the data was analyzed using Flow-Jo software.

**Cell Migration Assay**

Cell migration assay was performed as previously described (*6, 22*). Trans-well chambers (8 μm pore, Corning) were pre-equilibrated with serum-free media. CRL9855 monocytes (2 x

$10^4$/chamber) that were pre-stained for 1 hr per manufacturer's instructions with Far Red Cell Trace dye (Thermo Fischer Scientific), were placed in the top chamber, and 700 µl of control or FXR1 overexpression cells were placed in the bottom chamber. The chambers were incubated at 37°C for 18 hr in 5% $CO_2$. Cells on the upper surface of the filter were removed with a cotton swab. Migrated stained monocyte cells were observed in the bottom chamber and visualized using a microscope. Microscope images were taken, and the numbers of migrated cells were determined.

**Inhibitors**

Cytarabine (AraC) (*23, 24*), Trans-ISRIB (*25, 26*), FX-1 (*27-29*), and Toyocamycin (*30-34*) were obtained from Cayman Chemicals. FXR1 OE and control cells were treated with 1 µM trans-ISRIB for 24 h. For cell viability that was measured by Trypan blue staining and cell counts (*35*), FXR1 OE and control cells were treated individually or with a combination of 5 µM AraC and 1 µM trans-ISRIB for 24 h; 100 nM Toyocamycin and 1 µM AraC; 10 µM FX-1 and 5 µM AraC.

**Motif, GO, GSEA, and RNAFOLD analysis**

Multiple Em for Motif Elicitation (MEME) software was used to search for 5′UTR elements as described earlier (*35, 36*), using 5' UTR sequences from Genbank. Gene ontology (GO) analysis for differentially expressed translatome or proteome was conducted by DAVID 6.7 tools (*11*) (*12*) as described earlier (*35*) with our datasets. Gene Set Enrichment Analysis (GSEA) (*37, 38*) was performed as described earlier (*35*) with our datasets. 5′UTR folding as shown in Fig. S4A was performed using RNAFold web server from the Vienna RNA package (*39*) (rna.tbi.univie.ac.at/cgi-bin/RNAWebSuite/RNAfold.cgi).

**Statistical analyses**

Each experiment was repeated at least 3 times. No statistical method was used to pre-determine sample size. Sample sizes were estimated on the basis of availability and previous experiments (*2, 3*). No samples were excluded from analyses. P values and statistical tests were conducted for each figure. Statistical analyses were conducted using Excel. SEM (standard error of mean) values are shown as error bars in all figures. Means were used as center values in box plots. P-values less than 0.05 were indicated with an asterisk. E-values were used for the statistical significance in the motif analysis.

**Data Availability**

Raw datasets will be submitted to GEO public repository at final submission and will be available publicly as well as on request to the authors.

**Supplementary Table Legends:**

**Table S1.**

**a.** Table of RNA changes in G0 and serum-grown THP1 cells, with and without FXR1 depletion from (*6*).

**b**. SnoRNAs regulated in FXR1 KD G0 cells compared to control cells from sheet a.

**c.** SnoRNAs regulated in G0 cells compared to S+ cells from sheet a.

**d.** Common SnoRNAs derived from sheets b. and c.

**e.** Table of FXR1-RNA interactome. FXR1 immunoprecipitation followed by microarray from in vivo formaldehyde crosslinked THP1 cells grown under serum starved condition for 2 days to induce G0.

**f.** SnoRNAs bound to FXR1 from sheet e.

**g.** Table of rRNA modifications in FXR1 KD cells. rRNA enriched RNA fraction of FXR1 KD and control cells grown under serum starved condition for 1 day to induce G0, followed by LC MS/MS analysis of RNA nucleosides (ArrayStar service).

**Table S2**

Table of FXR1-protein interactome. FXR1 immunoprecipitation followed by TMT spectrometry from in vivo crosslinked THP1 cells grown under serum starved conditions for two days (2DG0) to induce G0.

**Table S3**

**a.** Table of TMT spectrometry analysis of protein levels in THP1 cells grown for four days under serum starved conditions to induce G0 (4DG0), and in THP1 cells treated with 5 μM AraC for 3 days (AraC), compared to untreated THP1 cells grown in medium with serum (S+) from our past dataset (*35*).

**b.** Ribosomal protein levels in THP1 cells grown for four days under serum starved conditions (4DG0) to induce G0, compared to THP1 cells grown in media with serum (S+), collated from sheet a.

**c.** RNA levels of ribosomal protein gene levels in FXR1 KD G0 cells (serum-starved for 1 day to induce G0) compared to control shRNA G0 cells, collated from Table S1 sheet a (from microarray analysis of mRNA level changes in FXR1 KD with 1 day G0 serum-starvation compared to control shRNA G0 cells from (*6*).

**d**. Microarray analysis of mRNA level changes in FXR1 KD 2 day G0 (serum-starved for 2 days to induce G0) and control shRNA cells from (*3*).

**e.** RNA levels of ribosomal protein genes in FXR1 KD 2 day G0 cells over control shRNA cells, collated from sheet d. RNA profiles were found to be comparable between 2 day G0 and 1 day G0 data.

**f.** TMT-spectrometry analysis of the proteome in FXR1 KD cells and control cells grown in serum or under serum starved conditions for 1 day (1DG0).

**g.** Level of ribosomal proteins in FXR1 KD cells compared control cells (proteome), collated from f.

**h.** Table of polysome associated mRNA in FXR1 KD G0 cells and control shRNA G0 cells (translatome by microarray analysis).

**i.** mRNA levels of ribosomal proteins on polysomes of FXR1 KD G0 cells over control shRNA G0 cells, collated from sheet h.

**Table S4**

**a.** Translationally regulated genes from the FXR1 KD and control datasets (Table S3 sheet h) whose mRNA levels were analyzed on polysomes of FXR1 OE cells by qPCR in Fig 4B.

**b.** Table of polysome associated mRNAs in FXR1 OE cells compared to control cells (translatome).

**c.** GSEA analysis of the translatome data in sheet b.

**d.** Overlap of increased translatome in FXR1 OE versus control cells in sheet b, and in AraC-treated cell versus untreated cell proteome from Table S3a.

**e.** Immune genes increased in the overlap of commonly increased genes in FXR1 OE translatome in sheet b, and in AraC-treated cell proteome from sheet d.

**f.** DAVID GO analysis of the downregulated FXR1 OE translatome (-1.5 fold and below from sheet b), showing genes in the category of phagocytosis inflammatory chemokine genes as well as immune receptor genes. Below, complete DAVID analysis of all downregulated (-1.5 fold and below from sheet b) gene set from the FXR1 OE translatome compared to control vector translatome. Below, GSEA analysis of the overlap of genes from all downregulated (-1.5 and below from sheet b) gene set from the FXR1 OE translatome compared to control vector translatome and from AraC-treated cell proteome from Table S3a.

**Table S5**

Oligonucleotides and shRNA target sequences used in the study.

**Supplementary Figure Legends:**

**Fig. S1.**

**A.** Western blot of FXR1 levels with actin as control in THP1 cells, untreated or treated with 1 μM and 5μM AraC for the indicated amount of time.

**B.** Western blot of FXR1 levels with actin as control in NOMO1 cells, untreated or treated with 1 μM and 5 μM AraC for the indicated amount of time. (See also Fig. 1 and Table S1)

**Fig. S2.**

**A.** Protocol flowchart illustrating enrichment of rRNA from FXR1 knockdown (FXR1 KD) and control THP1 G0 cells for LC-MS analysis of RNA modifications (top). Below table showing RNA mass spectrometry analysis of the 2′-*O*-methylated forms of adenosine, cytidine, guanosine, uridine, pseudo-uridylated form of uridine, and N4-acetylated form of cytidine compared to their respective unmodified forms. Modification analysis was done by Arraystar inc, where RNA was hydrolyzed to nucleosides before LC-mass spectrometry (Table S1g for complete data).

**B. a.** Top translation factors that co-immunoprecipitate with FXR1 in G0 THP1 cells. FXR1 was immunoprecipitated from in vivo crosslinked G0 THP1 cells followed by Tandem-Mass-Tag (TMT) spectrometry(*3, 14*) analysis to identify interacting proteins (Table S2 for complete list).

**b.** Western blot of FXR1 immunoprecipitation from the cytoplasmic fraction of in vivo crosslinked FXR1 overexpression (FXR1 OE) and control cells, followed by RT-qPCR for levels of 28S, 5S rRNA associated with FXR1 in Control and FXR1 OE cells. Shown qPCR of rRNA levels immunoprecipitated with FXR1 antibody normalized to IgG control.

**C.** TMT spectrometry proteomic analysis (from (*35*) shown in Table S3a in G0 and AraC-treated cells compared to untreated THP1 cells) revealing levels of POLR1D iso-2 in AraC treated and G0 cells compared to serum grown, untreated THP1 cells. (See also Fig. 2 and Tables S2, S3)

**Fig. S3.**

**A.** Diethylaminoethanol cellulose (DEAE) fractionation of in vivo crosslinked cytoplasmic extracts, reveals ribosome migration in G0, AraC-treated, and FXR1 OE THP1 cells compared to untreated S+ THP1 cells (migrating in increasing salt fractions). Shown qPCR analysis of both 28S and 18S rRNA levels (representing 80S ribosomes) in various fractions normalized to their levels in the inputs.

**B. a.** Protocol steps of experiment in Figure 3C. Mature ribosomes were immunoprecipitated with Y10B antibody from cytoplasmic fractions of in vivo crosslinked cells, followed by DEAE fractionation using increasing salt concentrations (top left). **b.** Western blot following Y10B immunoprecipitation showing RPLP0 verifying ribosome pull down (bottom left). Levels of 45S, and 5.8S rRNAs after immunoprecipitation with Y10B antibody from FXR1 OE and control cells. Shown qPCR of 45S and 5.8S rRNA verifying pull down of mature ribosome with 5.8S rRNA but not pre-ribosomal 45S rRNA (bottom right). **c.** Immunoprecipitation of ribosomes with Y10B antibody is verified by association with 5.8S rRNA. Shown qPCR of 5.8S rRNA level after immunoprecipitation with Y10B antibody in S+, G0, and AraC THP1 cells normalized to input levels (top right).

**C.** Fold RPLP0 (mRNA levels-total, polysome association levels, and protein levels from (*35*)) levels in FXR1 KD G0 cells compared to control cells.

**D.** Overexpression of snoRNAs in THP1 cells. Shown qPCR of snoRA2A and snoRD46 levels in THP1 cells after nucleofection with plasmids overexpressing the respective snoRNAs (Fig. 3D).

**E.** Western Blot of BT549 breast cancer cell lysates with control and FXR1 shRNAs showing levels of eIF2α phosphorylation upon FXR1 depletion.

**F.** RNA levels by qPCR of GUG reporter, AUG reporter normalized to tRNA-met and Renilla, in G0, AraC-treated, and FXR1 OE cells, for Luciferase reporter assays in Fig. 3G. (See also Fig. 3 and Table S3)

**Fig. S4.**

**A a**. Secondary structures and corresponding ΔG values of the 5′UTRs of top translated genes from the upregulated gene set translatome of FXR1 OE cells as predicted by the RNAFold web server (http://rna.tbi.univie.ac.at/cgi-bin/RNAWebSuite/RNAfold.cgi).

**b.** Enriched motif present in the 5'UTRs of the common set of genes that are increased at the protein level in THP1(G0), THP1 (AraC) cells and increased polysome association in FXR1 OE cells (generated by MEME analysis, from Table S4a-b) and Enriched motif present in the 5'UTRs of the genes with increased polysome association in FXR1 OE cells (generated by MEME analysis, from Table S4b).

**B a.** Table of known non-AUG bearing mRNAs (*40*) that show increased polysome association in FXR1 OE cells in the profiling dataset in Table S4b.

**b.** Enriched motif present in the 5'UTRs of the genes with increased polysome association in FXR1 OE cells and with known non-AUGs (generated by MEME analysis from the table in Ba.).

**C. a.** Microscope imaging of immune cell migration in a trans-well assay using Far-red stained monocytes on the top chamber, with either vector control cells or FXR1 OE cells in the bottom chamber, quantitated by imaging with graph shown in Fig. 4Cb.

**b.** Table of flow cytometry data showing survival of control vector cells compared to FXR1 OE cells (stained with Far-red cell trace dye as well as Hoechst 33342 to detect live cells) after co-culturing with monocytes and macrophages that were polarized or not polarized (M0 shown in Fig. 4Cc).

**D.** Model for FXR1-mediated ribosome alterations that induce translation changes to promote AML survival.
